# The FIGNL1-FIRRM complex is required to complete meiotic recombination in the mouse and prevents massive DNA damage-independent RAD51 and DMC1 loading

**DOI:** 10.1101/2023.05.17.541096

**Authors:** Akbar Zainu, Pauline Dupaigne, Soumya Bouchouika, Julien Cau, Julie A. J. Clément, Pauline Auffret, Virginie Ropars, Jean-Baptiste Charbonnier, Bernard de Massy, Raphael Mercier, Rajeev Kumar, Frédéric Baudat

## Abstract

During meiosis, nucleoprotein filaments of the strand exchange proteins RAD51 and DMC1 are crucial for repairing SPO11-generated DNA double-strand breaks (DSBs) by homologous recombination (HR). A balanced activity of positive and negative RAD51/DMC1 regulators ensures proper recombination. Fidgetin-like 1 (FIGNL1) was previously shown to negatively regulate RAD51 in human cells. However, FIGNL1’s role during meiotic recombination in mammals remains unknown. Here, we deciphered the meiotic functions of FIGNL1 and FIGNL1 Interacting Regulator of Recombination and Mitosis (FIRRM) using male germline-specific conditional knock-out (cKO) mouse models. Both FIGNL1 and FIRRM are required for completing meiotic prophase in mouse spermatocytes. Despite efficient recruitment of DMC1 on ssDNA at meiotic DSB hotspots, the formation of late recombination intermediates is defective in *Firrm* cKO and *Fignl1* cKO spermatocytes. Moreover, the FIGNL1-FIRRM complex limits RAD51 and DMC1 accumulation on intact chromatin, independently from the formation of SPO11-catalyzed DSBs. Purified human FIGNL1ΔN alters the RAD51/DMC1 nucleoprotein filament structure and inhibits strand invasion *in vitro*. Thus, this complex might regulate RAD51 and DMC1 association at sites of meiotic DSBs to promote proficient strand invasion and processing of recombination intermediates.

## Introduction

Meiosis ensures the accurate reduction of chromosome numbers in gametes during sexual reproduction. Erroneous meiosis results in sterility or fertility defects owing to aberrant gamete formation. During meiosis, homologous chromosomes (homologs) undergo pairing, synapsis, and recombination. Homologous recombination (HR) is crucial for crossover (CO) formation between homologs to ensure their balanced segregation during meiosis, and for promoting pairing and synapsis of homologs in some organisms including mammals ^1–3^. HR is initiated by genome-wide SPO11-dependent DNA double-strand break (DSB) formation ^4^. SPO11 is subsequently released from DSB sites as SPO11-oligonucleotide complex by resection machinery giving rise to 3’ single-stranded DNA (ssDNA) overhangs ^5,6^. The heterotrimeric complex of Replication Protein A (RPA) binds to and protects the ssDNA overhangs from nucleolytic degradation. Two eukaryotic RecA-like strand-exchange proteins, RAD51 and its meiosis-specific paralog DMC1, replace RPA on ssDNA with the help of the mediator protein BRCA2 ^7,8^. Both strand exchange proteins can catalyze homology search and strand exchange through invasion on an intact template, leading to formation of a joint molecule termed displacement loop (D-loop) ^9^. The invading end primes DNA synthesis that requires the dissociation of RAD51/DMC1 from double-strand DNA (dsDNA) within the D-loop. After D-loop formation, meiotic DSB repair can produce a non-crossover (NCO), or a CO by two alternative pathways that coexist in many organisms ^2^. In mice, the meiosis-specific class I CO pathway generates 90% of COs and is dependent on a set of proteins referred to as ZMM proteins (including the MSH4-MSH5 complex and TEX11) ^10,11^ and the MutL homologs MLH1-MLH3. Mouse MSH4 and MSH5 are essential to repair most if not all meiotic DSBs ^2,12–14^. The class II COs (∼10% of COs in the mouse) depend on structure-specific endonucleases ^2^.

Both RAD51 and DMC1 form foci, which colocalize extensively at DSB sites ^15,16^ and are proposed to assemble into side-by-side homo-filaments on ssDNA tails, with RAD51 at the DSB-distal region and DMC1 polymerizing on the 3’, DSB-proximal region ^9,17^. DMC1 is likely the main catalyzer of meiotic interhomolog recombination in most eukaryotes, while RAD51 plays crucial non-catalytic accessory roles ^18–20^. RAD51 is the sole strand exchange protein during mitotic recombination and also plays a strand exchange activity-independent role in the replication fork protection that might rely on its dsDNA-binding capacity ^21–26^. Besides this specific function, inactive filaments of RAD51 and DMC1 on dsDNA are likely toxic and are actively prevented ^27^. Members of the Swi2/Snf2-related RAD54 translocase family ^28^ prevent the accumulation of RAD51 on dsDNA in human cells ^29^, and of Rad51 and Dmc1 in *S. cerevisiae* ^30,31^. In *S. cerevisiae*, Rad54 and its paralog Rdh54 promote strand invasion, and remove RAD51/DMC1 from dsDNA following D-loop formation ^28,32^. In mouse, RAD54 and its paralog RAD54B are not essential for meiotic recombination, because the *Rad54 Rad54b* double mutant mice are fertile ^33,34^. Many proteins regulate RAD51/DMC1 nucleofilament formation positively or negatively. Positive factors are required to form stable and active RAD51-ssDNA filaments. They include BRCA2 and several RAD51 paralogs, which are essential for viability in mammals ^7,8^. The conserved Shu complex comprises in mammals the distant RAD51 paralog SWSAP1, the SWIM-domain containing SWS1 and SPIDR ^35–39^. This complex promotes RAD51-mediated HR in the context of replication, and RAD51- and DMC1-mediated meiotic recombination in the mouse, but is not essential for viability ^36,37,40–42^. The SWSAP1-SWS1-SPIDR complex might promote specifically the stable assembly of longer RAD51 nucleoprotein filaments involved in some HR types, especially interhomolog HR ^36,37^.

FIGNL1 (fidgetin-like 1) forms an evolutionary conserved complex with FIRRM (FIGNL1 interacting regulator of recombination and mitosis) that interacts with RAD51 and DMC1, and regulates negatively RAD51 during HR repair in mammalian cells ^43–51^. In Arabidopsis and rice meiosis, FIGNL1 and FIRRM homologs negatively regulate the dynamics of RAD51 and DMC1 foci and limit the formation of class II crossovers ^44,45,52–54^. Arabidopsis *figl1* (*Figl1* homolog) and *flip* (*Firrm* homolog) mutants are fertile with all meiotic DSBs repaired ^44,52,55^. Conversely, unrepaired DSBs persist in rice *fignl1* and *meica* (*Firrm* homolog) mutants, leading to chromosome fragmentation and sterility ^53,54^. The regulation of RAD51/DMC1 focus formation in Arabidopsis somatic and meiotic cells involves an antagonistic interplay between BRCA2 and FIGL1, consistent with FIGL1 acting as a negative regulator of RAD51/DMC1 filament ^56^. In human cells, an antagonistic relationship was found between BRCA2 and FIGNL1-BRCA2 ^48^, and between the SWSAP1-SWS1-SPIDR complex and FIGNL1, which interacts with SWSAP1 ^40^ and SPIDR ^43^. Indeed, FIGNL1 depletion relieves the dependency on SWSAP1 and SWS1 for forming RAD51 repair foci ^40^. Moreover, purified human SWSAP1 protects RAD51-ssDNA filament from dissociation promoted by FIGNL1 *in vitro* ^40^. However, the role of FIGNL1 and FIRRM during meiotic recombination in mammals was unknown.

In this study, we investigated the role of the FIGNL1-FIRRM complex in meiotic recombination by analyzing germ line-specific mouse conditional knock-out models for both genes. The depletion of FIGNL1 or FIRRM in mouse spermatocytes results in meiotic DSB repair failure and no full synapsis between homologs during meiotic prophase I, leading to prophase I arrest and apoptosis. Surprisingly, *Fignl1* cKO and *Firrm* cKO spermatocytes also show an abundant DSB-independent accumulation of RAD51 and DMC1 on chromatin and meiotic chromosome axes during premeiotic replication and early meiotic prophase stages. This indicates that the FIGNL1-FIRRM complex prevents the formation of stable inactive RAD51 and DMC1 filaments, presumably on intact dsDNA, in mouse spermatocyte nuclei.

## Results

### FIGNL1 and FIRRM are required for meiotic prophase completion in the mouse male germline

We wanted to determine the roles of FIGNL1 and its putative partner FIRRM (also called BC055324) during meiosis. As both genes are essential for mouse viability (^57,58^, IMPC, https://www.mousephenotype.org/), we generated cKO lines with Cre expression under the control of the *Stra8* promoter ^59^ to ablate *Firrm* or/and *Fignl1* in the male germline shortly before meiosis onset (*Firrm* cKO and *Fignl1* cKO, Extended Data Fig. 1a-c). Testis weight was similarly reduced in *Firrm* cKO, *Fignl1* cKO, and *Firrm-Fignl1* double cKO mice compared with wild-type controls (Fig. 1a, Extended Data Fig. 2a). Analysis of testis sections from adult *Firrm* cKO and *Fignl1* cKO animals revealed the lack of haploid cells (spermatids) in most tubules, suggesting a prophase I arrest (Fig. 1b, Extended Data Fig. 2b-c). Some germ cell depletion was apparent on sections from older *Firrm* cKO and *Fignl1* cKO animals, suggesting that FIGNL1-FIRRM depletion affects the viability or the proliferative capacity of spermatogonia in adults (Extended Data Fig. 2b-c). Nevertheless, the density of Sertoli cells, spermatogonia and early prophase spermatocytes appeared normal in seminiferous tubules from 4-week-old *Firrm* cKO animals (Fig. 1b), and no obvious defect was detected in spermatogonia by cytological analysis of juvenile testes, suggesting that the defects observed in *Firrm* cKO and *Fignl1* cKO spermatocytes do not result from defects in spermatogonia prior to meiosis induction. The presence of some tubules with a small number of round spermatids and of few tubules with many round and elongated spermatids suggested incomplete or delayed Cre-mediated excision or protein depletion, as described in other conditional mouse mutants obtained with this *Stra8-Cre* transgene ^60–62^. Consistent with delayed excision or protein depletion, no sperm was found in cauda epididymis from 4-month-old mice (Extended Data Fig. 2d), and CRE-mediated *Firrm* excision was 100% efficient, as assessed by genotyping the progeny of *Firrm^flox/+^ Stra8-Cre^Tg^* males crossed with wild-type females (Methods). In testes from 12-day post-partum (12 dpp) *Firrm* cKO and *Fignl1* cKO mice, FIRRM and FIGNL1 protein levels in the cytoplasmic and nuclear fractions were greatly and similarly reduced compared to controls (Fig. 1c, Extended Data Fig. 2e). This suggests that they reciprocally regulate their stability, which is consistent with forming a complex. The residual protein level might result from expression in non-meiotic cells (spermatogonia or somatic cells) and/or from incomplete Cre-induced gene deletion in a fraction of spermatocytes. Conversely, RAD51 expression in the nuclear fraction was increased in *Firrm* cKO and *Fignl1* cKO testes (Fig. 1c, Extended Data Fig. 2e), suggesting that the FIGNL1-FIRRM complex might be implicated in limiting directly or indirectly nuclear accumulation of RAD51 (but not of DMC1). This might have significant consequences, because RAD51 nuclear level is suggested to play a role in HR regulation ^63^.

**Figure 1.**
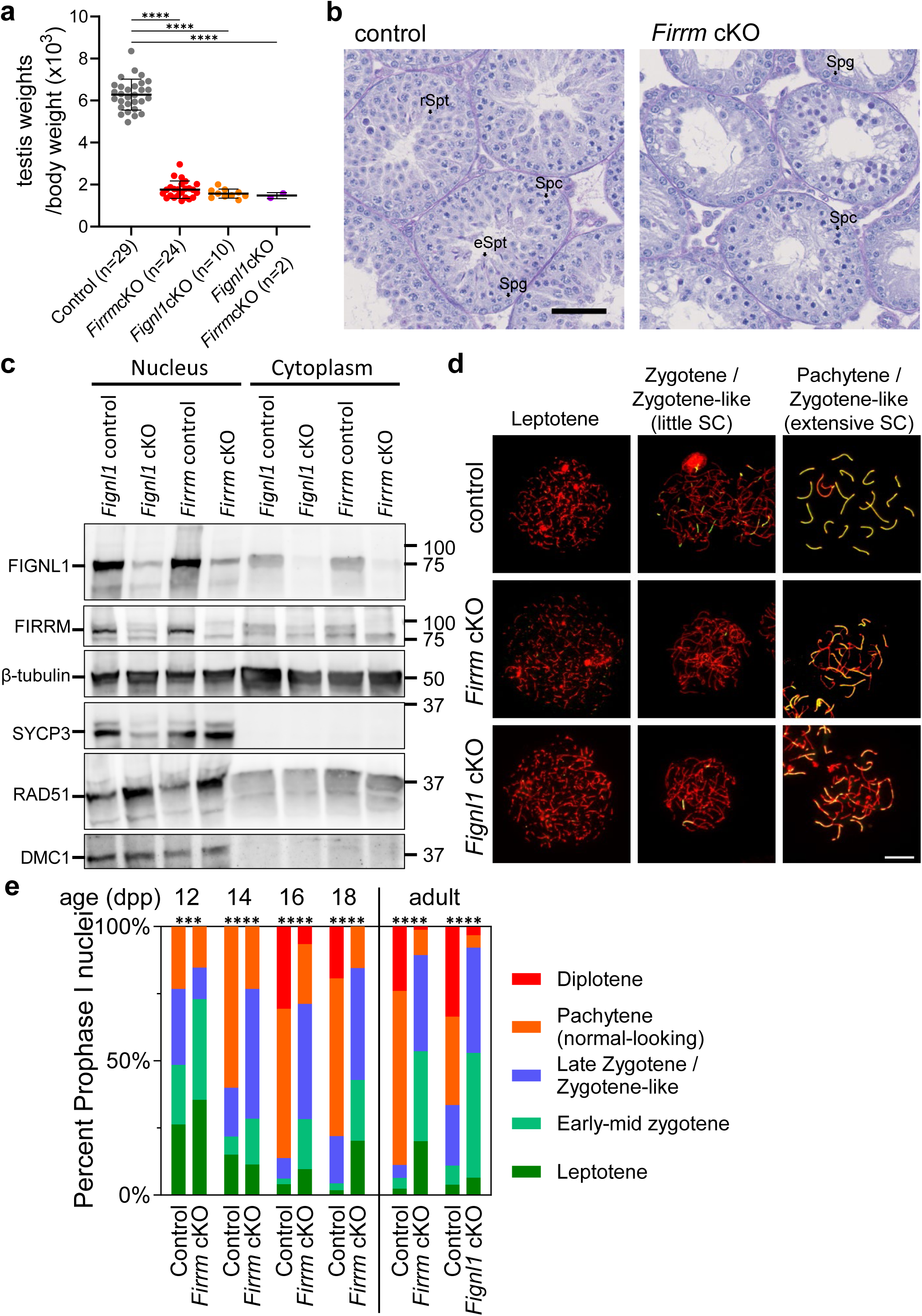
Spermatogenesis is defective in *Firrm* cKO and *Fignl1* cKO mice. **a.** Testis weight relative to body weight in control (n=29), *Firrm* cKO (n=24), *Fignl1* cKO (n=10) and *Firrm* cKO *Fignl1* cKO (n=2) adult mice (30 dpp to 154 dpp; kinetics is shown in Extended Data Fig. 2a). Unpaired t-test, two-sided. **b.** Periodic acid-Schiff-stained testis sections from 4-week-old control and *Firrm* cKO mice. Spg, spermatogonia; Spc, spermatocytes; rSpt, round spermatids; eSpt, elongated spermatids. Scale bar, 50 µm. **c.** Western blot analysis of cytoplasmic (80µg) and nuclear (100µg) fractions from testes of 12 dpp mice of the indicated genotypes. **d.** Chromosome axes (SYCP3, red) and synaptonemal complex (SYCP1, green) were detected in spread spermatocyte nuclei from control, *Firrm* cKO (25 dpp) and *Fignl1* cKO (56 dpp) mice. Scale bar, 10 µm. **e.** Distribution of spermatocytes at different meiotic prophase substages in juvenile *Firrm* cKO mice (indicated age) and in adult (8-week-old) *Firrm* cKO and *Fignl1* cKO mice. Chi-square test. For all figures: ns, non-significant; *0.01< p ≤0.05; **0.001< p ≤0.01; ***0.0001< p ≤0.001; ****p ≤0.0001.

The synaptonemal complex (SC), a tripartite proteinaceous structure, links the axes of homologous chromosomes during meiotic prophase. To analyze if synapsis was impaired in *Firrm* cKO and *Fignl1* cKO, we stained surface-spread spermatocyte nuclei with antibodies against SYCP3, a component of meiotic chromosome axes, and SYCP1, a protein of the SC central element (Fig. 1d). *Firrm* cKO and *Fignl1* cKO spermatocytes formed apparently normal meiotic chromosome axes (leptotene stage), suggesting a normal meiotic prophase entry. However, most nuclei showed unsynapsed or partially synapsed axes, indicating the accumulation of zygotene-like cells. The small fraction of *Fignl1* cKO and *Firrm* cKO spermatocytes that progressed toward normal-looking pachytene and diplotene might be explained by incomplete Cre-mediated excision in these cells. We followed prophase I progression during the first wave of meiosis in *Firrm* cKO, from 12 dpp to 18 dpp (Fig. 1e). We detected a deficit in more advanced stages already in 12 dpp *Firrm* cKO spermatocytes. At 16 and 18 dpp, most nuclei were arrested at a zygotene-like stage, and the percentage of nuclei at the pachytene and diplotene stages was strongly reduced (at 18 dpp, 78% of control versus 15% of *Firrm* cKO nuclei). Approximately 30% of *Firrm* cKO prophase I nuclei displayed an abnormal zygotene/pachytene-like pattern, with non-homologous synapsis and only few synapsed chromosome axes (Fig. 1d). These findings in 12 dpp to 18 dpp spermatocytes are indicative of an arrest in early pachytene and a defect in homologous synapsis. Adult *Firrm* cKO and *Fignl1* cKO animals displayed a similar deficit in pachytene-diplotene spermatocytes (Fig. 1e), consistent with the hypothesis that FIGNL1 and FIRRM act together.

### The formation and initial processing of meiotic DSBs are unaffected in *Firrm* cKO and *Fignl1* cKO spermatocytes

A synapsis defect associated with a mid-prophase arrest might result from defective recombination initiation (e.g. *Spo11*^-/- 64,65^) or defective repair of meiotic DSBs (e.g. *Dmc1^-/-^* ^66,67^) ^3^. To determine whether DSB formation was altered by FIRRM or FIGNL1 depletion, we quantified phosphorylated H2AX (γH2AX) that decorates chromatin in a DSB-specific manner at leptotene ^68^. The γH2AX signal intensity in the nucleus was not different in control and *Firrm* cKO spermatocytes from pre-leptotene (stage of pre-meiotic replication) to mid-late leptotene (Fig. 2a-b). RPA2, a subunit of the ssDNA-binding protein complex Replication Protein A (RPA), which is involved in DNA replication and HR, forms multiple foci at replication forks in preleptotene spermatocytes, and along chromosome axes at sites of recombination intermediates from leptotene to pachytene stage ^60,69,70^. RPA2 foci displayed a similar kinetics in *Firrm* cKO, *Fignl1* cKO, and control spermatocytes (Fig. 2c-d). An increase in zygotene spermatocytes (by 1.3-fold for both *Firrm* cKO and *Fignl1* cKO in early zygotene stage) might result from a different kinetics of DSB formation, from a difference in the formation or stability of HR intermediates involving detectable RPA, or from a difference in staging the nuclei, due to less or delayed synapsis initiation in mutant nuclei. Thus, the first steps of meiotic recombination (DSB formation and RPA recruitment on resected ssDNA ends) were not affected by the absence of the FIGNL1-FIRRM complex.

**Figure 2.**
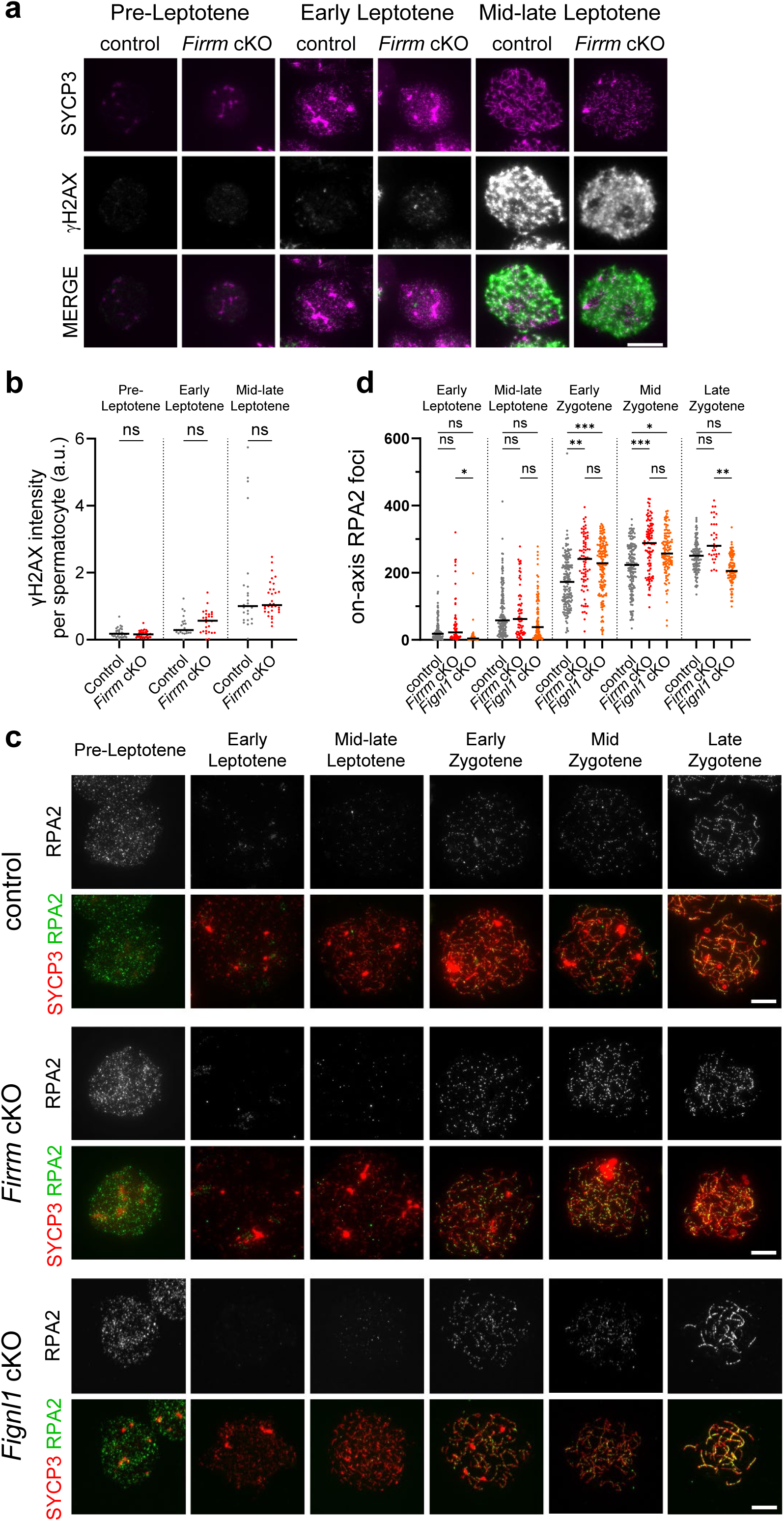
*Firrm* cKO and *Fignl1* cKO spermatocytes are proficient for markers of meiotic DSB formation and processing. **a.** Spread nuclei of spermatocytes from 12 dpp control and *Firrm* cKO mice stained for SYCP3 and γH2AX. Scale bar, 20 µm. **b.** Total nuclear γH2AX signal intensity in 12 dpp control (gray) and *Firrm* cKO (red) spermatocytes (n=2 mice per genotype). Unless stated otherwise, Dunn’s multiple comparison tests were performed in all focus number comparisons. **c.** Spread spermatocyte nuclei from 12 dpp control and *Firrm* cKO mice stained for SYCP3 and RPA2. Scale bar, 10 µm. **d.** Number of on-axis RPA2 foci in control (gray), *Firrm* cKO (red) and *Fignl1* cKO (orange) spermatocytes. control, n=6 (4x 12 dpp, 1x 16 dpp, 1x 17 dpp), *Firrm* cKO, n=4 (12 dpp) and *Fignl1* cKO, n=3 (1x 16 dpp, 2x 17 dpp) mice.

### The recruitment of RAD51 and DMC1 on meiotic chromatin strongly increases in the absence of FIGNL1 or FIRRM

In mouse spermatocytes, RAD51 and DMC1 foci extensively colocalize on meiotic chromosome axes (on-axis foci) from leptotene to pachytene stage, particularly in zygotene ^15,16,69^. Compared with controls, RAD51 and DMC1 signal intensity and foci pattern and kinetics were strikingly different in *Firrm* cKO and *Fignl1* cKO spermatocytes (Fig. 3a-c; Extended Data Fig. 3a). First, RAD51 (but not DMC1) formed many foci at preleptotene, during premeiotic replication. Second, the mean number of RAD51 and DMC1 on-axis foci was significantly higher in *Firrm* cKO and *Fignl1* cKO than in control spermatocytes at every stage, from early leptotene to zygotene. Third, in cKO spermatocytes, many RAD51 and DMC1 foci were located away from the chromosome axes (off-axis foci). The number of off-axis foci was highest during leptotene and progressively decreased during zygotene. Fourth, in cKO spermatocytes, RAD51 and DMC1 staining formed continuous lines, at our resolution limit, along the synaptonemal complex segments in zygotene-like nuclei. This did not allow counting RAD51 and DMC1 foci in late zygotene-like nuclei with extensive synapsis. Overall, these observations are consistent with the role of FIRRM and FIGNL1 in limiting RAD51 and DMC1 loading on chromatin in spermatocyte nuclei. We describe these different features in the following sections.

**Figure 3.**
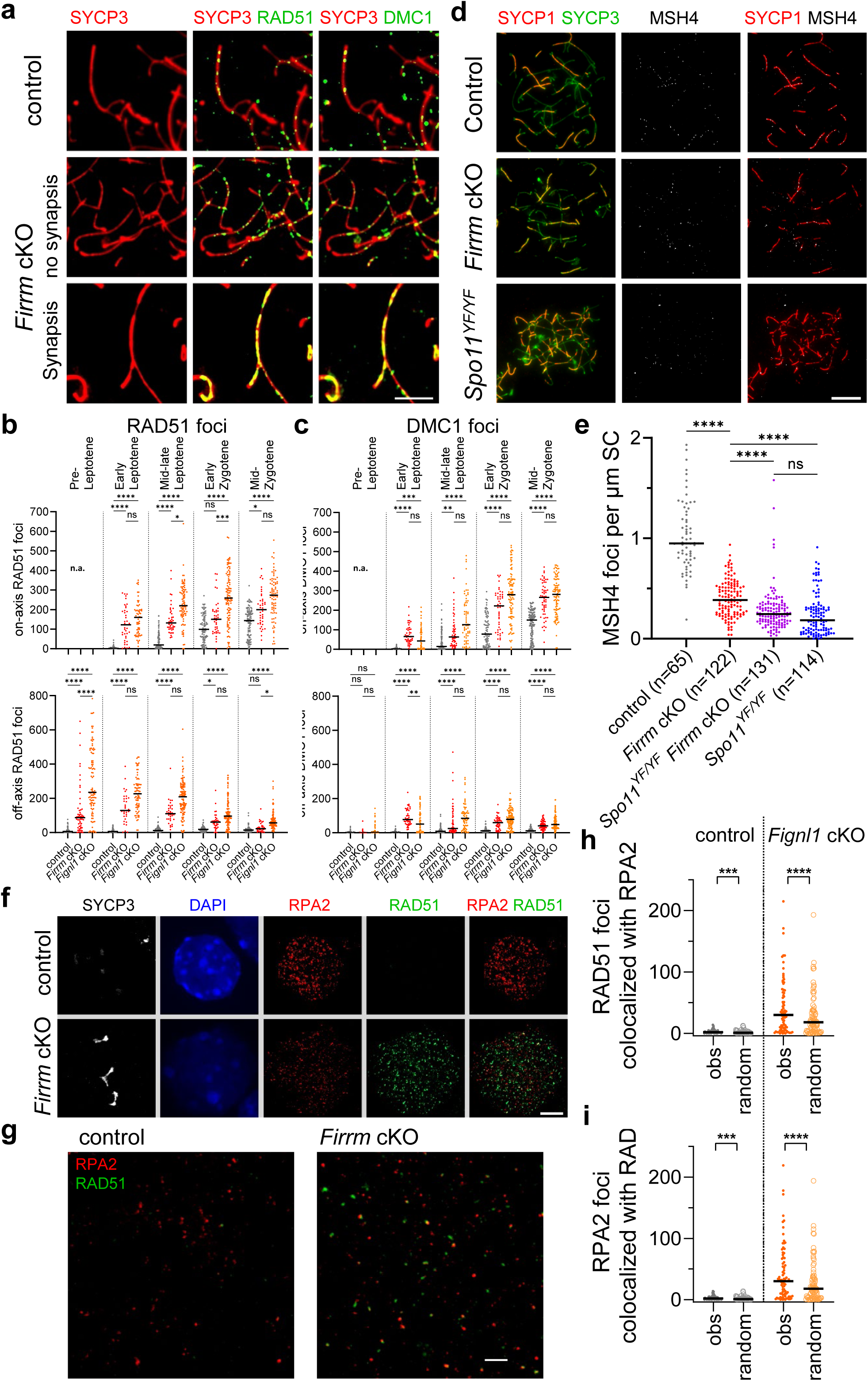
*Firrm* cKO and *Fignl1* cKO spermatocytes accumulate RAD51 and DMC1, but are defective for forming foci of proteins involved in late acting recombination proteins. **a.** Zygotene spermatocyte spreads from control and *Firrm* cKO mice stained for SYCP3, RAD51 and DMC1. Scale bar, 5 µm. **b, c.** Numbers of RAD51 **(b)** and DMC1 **(c)** foci in control, *Firrm* cKO and *Fignl1* cKO spermatocytes **(c)**. n=2 mice per genotype. **d.** Spreads of zygotene spermatocytes from 16 dpp control, *Firrm* cKO and *Spo11^YF/YF^* mice stained with SYCP3, SYCP1 and MSH4. Scale bar, 10 µm. **e.** MSH4 focus density along SYCP1-marked synaptonemal complex fragments in control, *Firrm* cKO, *Spo11^YF/YF^ Firrm* cKO, and *Spo11^YF/YF^* zygotene/zygotene-like spermatocytes. n=3 mice per genotype. **f.** Preleptotene spermatocyte spreads from control and *Firrm* cKO mice stained for SYCP3, RPA2 (red) and RAD51 (green). Scale bar, 10 µm. **g.** STED images of preleptotene spermatocyte spreads from control and *Firrm* cKO mice stained for RAD51 (STAR ORANGE, green) and RPA2 (STAR RED, red). Scale bar, 1 µm. **h-i.** Number of RAD51 foci that colocalized with RPA2 foci **(h)** and of RPA2 foci that colocalized with RAD51 **(i)** in spreads of preleptotene control and *Fignl1* cKO spermatocyte nuclei. n=2 (control) or n=3 mice (*Fignl1* cKO). The observed (obs) and expected by chance (random) numbers of colocalized foci are shown. Wilcoxon two-tailed test.

### Post-strand invasion recombination foci are strongly reduced in the absence of FIRRM

The efficient recruitment of RAD51 and DMC1 prompted us to examine MSH4 and TEX11, two meiotic stabilizing post-strand invasion recombination intermediate ZMM proteins ^11^, which form foci on SC from zygotene to mid-pachytene ^3,69,71^. The number of MSH4 and TEX11 foci was strongly reduced in late zygotene-like *Firrm* cKO nuclei compared with the control (Fig. 3d, Extended Data Fig. 3b). To normalize differences in SC extension among genotypes, we measured the density of MSH4 foci per µm of SC length. MSH4 focus density was reduced by 2.5-fold in *Firrm* cKO compared with control spermatocytes (Fig. 3e, Extended Data Fig. 3c), although the number of MSH4 foci was higher than in *Spo11^YF/YF^* nuclei (without DSB-inducing activity). Thus, FIRRM is required for TEX11 and MSH4 focus formation during mouse meiotic recombination. The residual MSH4 foci might result from MSH4 binding to a small fraction of normal or aberrant recombination intermediates formed in the absence of the FIGNL1-FIRRM complex. Alternatively, we cannot exclude the persistence of a small amount of FIRRM protein in *Firrm* cKO spermatocytes, sufficient for recruiting MSH4 to few recombination intermediates. Thus, despite the increased recruitment of RAD51 and DMC1 on chromosome axes, the processing of recombination intermediates was defective in *Firrm* cKO spermatocytes, suggesting a function of FIGNL1-FIRRM at a step likely before recombination intermediate stabilization by MSH4-MSH5.

### RAD51 accumulates on chromatin during premeiotic replication in *Firrm* cKO and *Fignl1* cKO spermatocytes

RPA2 forms many foci at ongoing replication forks in preleptotene nuclei (Fig. 3f-g). The kinetics of RPA2 focus formation was similar in control, *Firrm* cKO and *Fignl1* cKO spermatocytes, and few foci remained in early leptotene stage. This suggests that premeiotic replication was completed without major alteration (Fig. 2d; Extended Data Fig. 3d). As RAD51 is involved in protecting stalled replication forks ^72^, we hypothesized that RAD51 might colocalize with RPA during premeiotic replication in *Firrm* cKO and *Fignl1* cKO spermatocytes. We measured the colocalization of RAD51 and RPA2 in preleptotene spermatocytes and compared these data with the colocalization of randomly distributed foci obtained from simulations (see Methods; Fig. 3h-i; Extended Data Fig. 3e). In *Fignl1* cKO, 17% of RAD51 foci colocalized with RPA2 foci compared with 9% of randomly generated RAD51 foci (p <0.0001; two-tailed Wilcoxon test), suggesting that a fraction of RAD51 foci localizes at replication forks. However, the majority of identified RAD51 foci did not colocalize with RPA2 foci, suggesting that a larger fraction of RAD51 foci may not localize at replication forks. We cannot exclude that both RAD51 and RPA localize at forks in an exclusive manner, and that RAD51 binding excludes RPA. However, because of the large number of RAD51 foci that persisted at the end of premeiotic replication and the absence of obvious gross replication defects, we hypothesize that RAD51 colocalizes transiently with RPA at replication forks. It then remains in place, possibly on intact DNA, while the forks progress and move away. DMC1 foci were rare in most *Firrm* cKO and *Fignl1* cKO preleptotene spermatocytes (Fig. 3c), likely because meiosis-specific DMC1 production is still low at preleptotene stage.

### RAD51 and DMC1 form parallel linear structures along the synaptonemal complex in the absence of FIRRM

To refine the characterization of RAD51 and DMC1 distribution in *Firrm* cKO spermatocytes, we visualized RAD51, DMC1 and SYCP3 using super-resolution stimulated emission depletion (STED) microscopy (Fig. 4a-b). In leptotene and zygotene control spermatocytes, RAD51 and DMC1 formed partially overlapping co-foci along the unsynapsed axes and SC segments. RAD51 was more often closer to the chromosome axis than DMC1, as described previously ^20,73^. In *Firrm* cKO spermatocytes, the patterns of RAD51 and DMC1 staining were more heterogeneous. A first type of RAD51-DMC1 co-foci was similar to control foci, but RAD51 signal tended to be more extended compared with DMC1 (Fig. 4a, compare insets i-iv, control, with insets v-vi, *Firrm* cKO). Consistent with this observation, the intensity of RAD51 foci was higher in *Firrm* cKO than in control spermatocytes, whereas DMC1 focus intensity was unchanged or even lower (Extended Data Fig. 4a-b). Second, some RAD51-DMC1 co-foci formed longer structures anchored to the chromosome axis, a pattern expected if they were extending along chromatin fibers (Fig. 4a, inset vii). The localization patterns of these two types of foci are compatible with RAD51/DMC1 filaments bound to chromatin fibers at DSB sites or/and dsDNA. In addition, at some sites, RAD51 and DMC1 followed the unsynapsed axes, often filling gaps with little or no SYCP3 signal between more heavily SYCP3-stained axis segments (Fig. 4a, insets viii-ix). Lastly, in *Firrm* cKO zygotene-like nuclei with partial synapsis, RAD51 and DMC1 formed two parallel lines separated by ∼100 nm along SC segments, between the lateral elements (axes) visualized by ∼210 nm apart SYCP3 signal lines (Fig. 4b-c). The intensity of these lines was irregular with interruptions, and interspersed with more intense foci. These observations indicate a highly aberrant patterning of RAD51 and DMC1 on meiotic chromatin and on meiotic chromosome axes in the absence of FIGNL1 and FIRRM activity.

**Figure 4.**
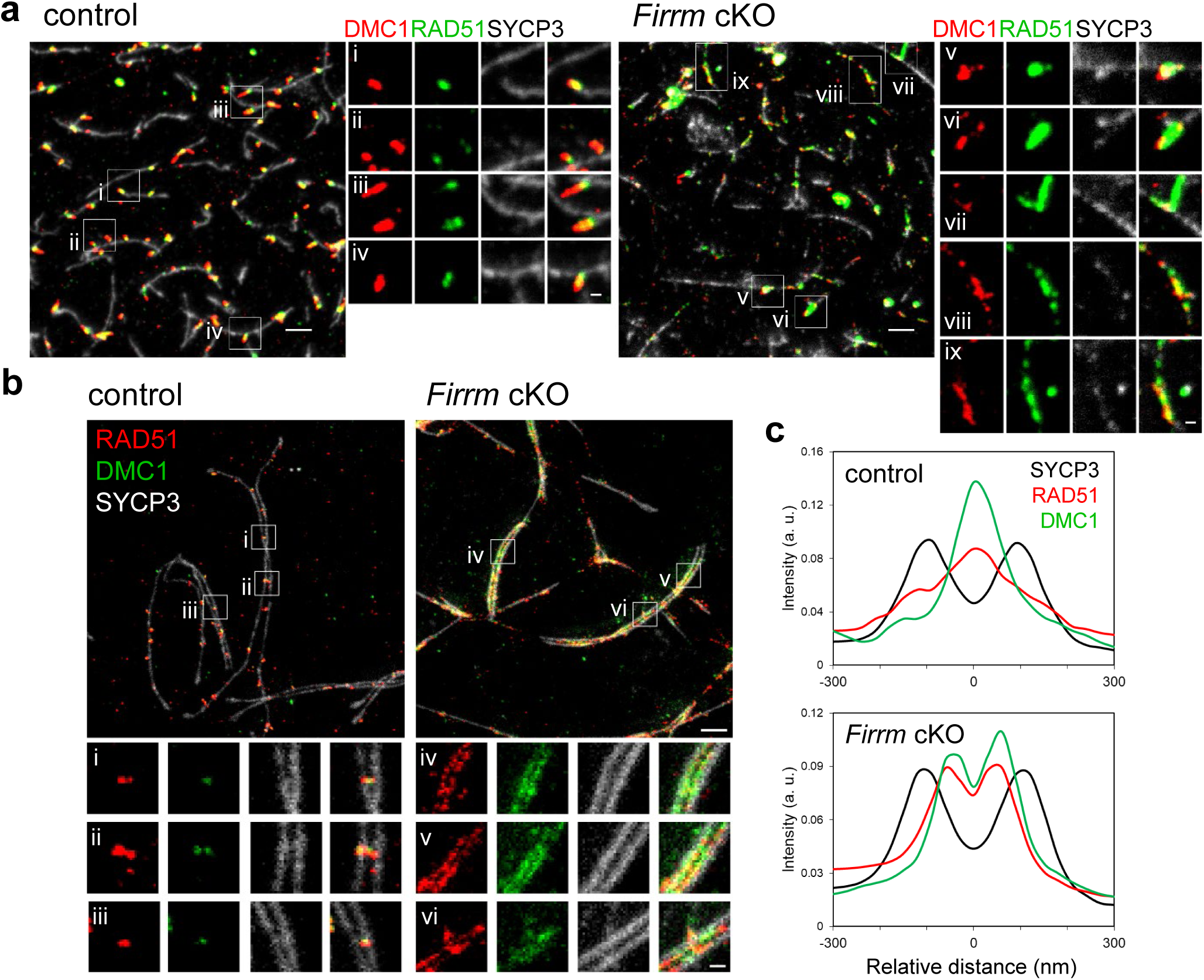
RAD51 and DMC1 patterns in mouse meiotic chromosomes. **a.** STED images of spreads of leptotene spermatocyte nuclei stained for SYCP3 (STAR GREEN, white), RAD51 (STAR ORANGE, green), and DMC1 (STAR RED, red). **b.** STED images of spreads of zygotene/zygotene-like spermatocyte nuclei with extensive synaptonemal complexes, stained for SYCP3 (STAR 460L, white), RAD51 (STAR RED, red) and DMC1 (STAR ORANGE, green). **c.** Relative intensity of SYCP3 (black), RAD51 (red) and DMC1 (green) signal across the synaptonemal complex in control (across RAD51-DMC1 mixed foci) and *Firrm* cKO (outside regions of stronger focus-like RAD51-DMC1 staining). Mean of 12 sections from STED images of 3 different nuclei.

### Accumulation of RAD51 and DMC1 foci in *Firrm* cKO spermatocytes is meiotic DSB-independent

In *Firrm* cKO and *Fignl1* cKO spermatocytes, RAD51 and DMC1 displayed an unusual pattern that included an increased number of foci, many off-axis foci, and linear staining along chromosome axes. This was different from the expected discrete DSB repair foci on chromosome axes ^16,69^, raising the question of whether in these cKO models, RAD51 and DMC1 recruitment requires SPO11-generated DSBs. Thus, we generated *Spo11^YF/YF^ Firrm* cKO double mutants in which SPO11 is catalytically dead ^74^. The low early prophase γH2AX staining and the absence of RPA2 foci confirmed the absence of DSBs in these animals (Extended Data Fig. 4c-d). Strikingly, we detected large numbers of on-axis and off-axis RAD51 and DMC1 foci in *Firrm* cKO and in *Spo11^YF/YF^ Firrm* cKO spermatocytes, and only background signal in *Spo11^YF/YF^*spermatocytes (as expected) (Fig. 5a). Overall, the pattern of RAD51 and DMC1 in *Firrm* cKO and *Spo11^YF/YF^ Firrm* cKO were quite similar: a large number of on-axis foci detected from early prophase that persisted through zygotene, RAD51 foci formed during preleptotene stage, and both RAD51 and DMC1 off-axis foci progressively disappeared from leptotene to zygotene (Fig. 5b-c; Extended Data Fig. 4e-f). RAD51 and DMC1 foci were consistently more numerous in single *Firrm* cKO than in *Spo11^YF/YF^ Firrm* cKO in early leptotene and zygotene, although the difference was on the merge of significance. In zygotene, the recruitment of RAD51 and DMC1 to SPO11-catalyzed DSB sites may explain this difference. It is more difficult to interpret in early leptotene, a stage with very few processed DSBs in *Firrm* cKO spermatocytes, based on the low number of RPA2 foci (Fig. 2d). Altogether, these observations demonstrate that the accumulation of RAD51 and DMC1 Firrm cKO spermatocytes is largely independent of SPO11-catalyzed DSBs.

**Figure 5.**
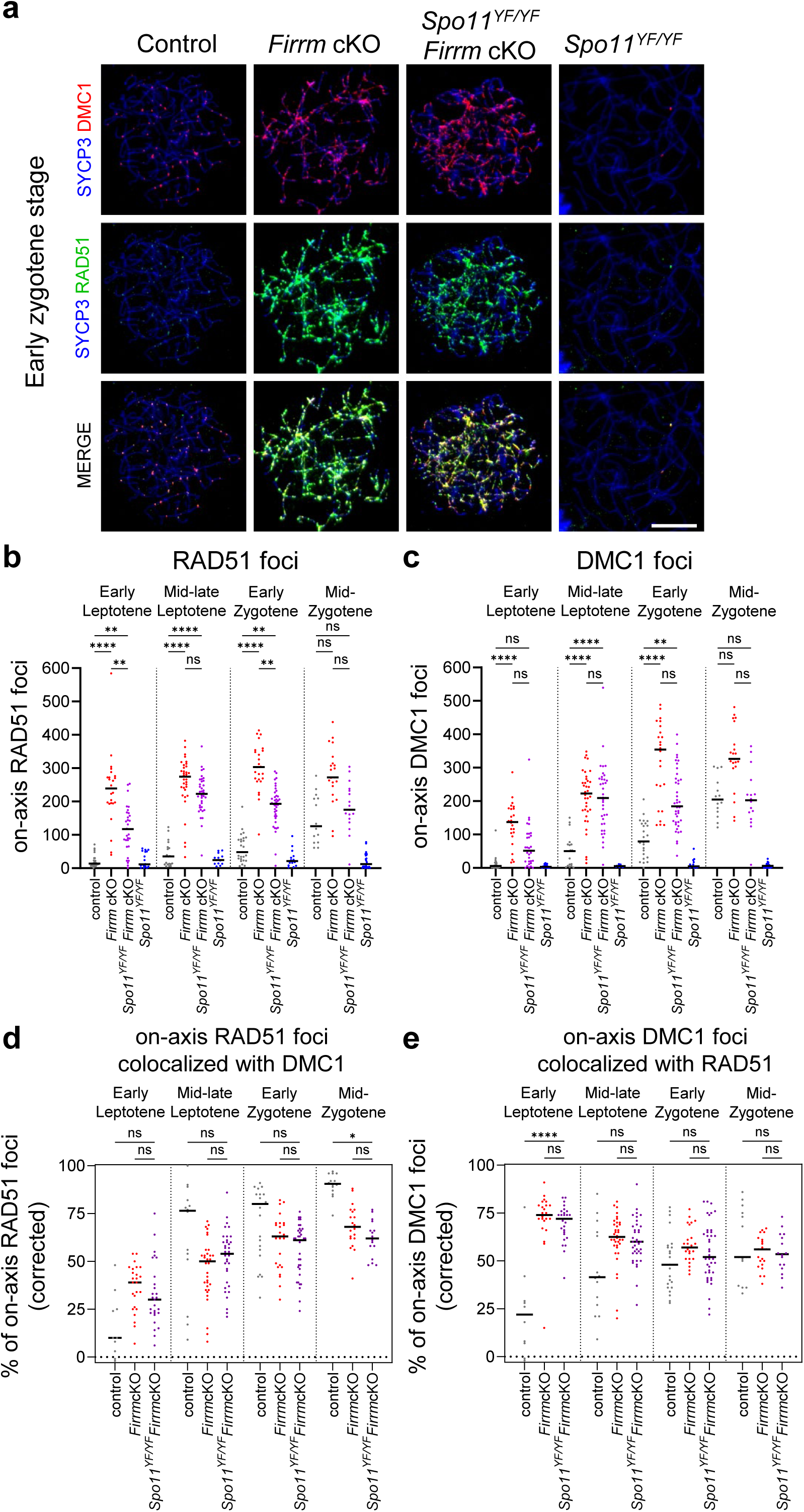
FIRRM prevents DSB-independent accumulation of RAD51 and DMC1 in mouse spermatocyte chromosomes. **a.** Spreads of representative control, *Firrm* cKO, *Spo11^YF/YF^ Firrm* cKO, and *Spo11^YF/YF^* early zygotene spermatocytes stained for SYCP3, DMC1 and RAD51. Scale bar, 10 µm. **b-c.** Counts of on-axis RAD51 **(b)** and DMC1 **(c)** foci in spreads from control, *Firrm* cKO, *Spo11^YF/YF^ Firrm* cKO, and *Spo11^YF/YF^*spermatocytes from 12 dpp mice. n=2 mice per genotype. **d-e.** Percentage (corrected for random colocalization, see Methods) of on-axis RAD51 foci colocalized with on-axis DMC1 foci **(d)** and vice-versa **(e)**, in spreads from control and *Firrm* cKO spermatocytes from 12 dpp mice.

In wild-type meiosis, RAD51 and DMC1 colocalization throughout the meiotic prophase reflects their cooperation at resected DSB ends ^15–17,20,73^. To determine their level of colocalization in *Firrm* cKO spermatocytes where RAD51 and DMC1 foci form in the absence of SPO11-catalyzed DSBs, we examined their colocalization from early leptotene to mid-zygotene stage, in nuclei containing at least 10 foci for each protein (Fig. 5d-e; Extended Data Fig. 5). As expected, on-axis RAD51 foci, the number of which was higher, colocalized less frequently with DMC1 foci in *Firrm* cKO than in control spermatocytes, especially at earlier stages (Fig. 5d). Conversely, on-axis DMC1 foci colocalized at least as frequently with on-axis RAD51 foci in *Firrm* cKO and *Spo11^YF/YF^ Firrm* cKO than in control spermatocytes at every stage, with a maximum in early leptotene (∼70%; Fig. 5e). Off-axis foci displayed the same trend, with a very high percentage of DMC1 foci that colocalized with RAD51 foci at earlier stages in *Firrm* cKO (Extended Data Fig. 5c-e). Moreover, RAD51 and DMC1 colocalization levels were similar in *Firrm* cKO and *Spo11^YF/YF^ Firrm* cKO, indicating that their association is DNA damage-independent. Altogether, these observations indicate that in the absence of FIRRM, off-and on-axis RAD51 foci assemble independently of DMC1 foci in preleptotene and early prophase spermatocytes. Detectable DMC1 foci might form by joining pre-existing RAD51 foci, or by co-assembling *de novo* RAD51-DMC1 foci in *Firrm* cKO spermatocytes.

### DMC1 is recruited to DSB sites in the absence of the FIGNL1-FIRRM complex

The abundance of DSB-independent RAD51 and DMC1 foci raises the question of whether these recombinases are recruited at meiotic DSB sites at all in the absence of FIRRM or FIGNL1. Therefore, we determined the colocalization of on-axis RAD51 or DMC1 foci with RPA2 foci, used as a marker of a subset of the DSBs, in *Firrm* cKO and *Fignl1* cKO spermatocytes. The number of on-axis RAD51-RPA2 and DMC1-RPA2 co-foci (corrected for random colocalization) in spermatocytes followed the kinetics of RPA2 foci in all genotypes (Extended Data Fig. 6a-c, compare with Fig. 2d). In *Firrm* cKO and *Fignl1* cKO spermatocytes, the percentage of on-axis RPA2 foci that colocalized with RAD51 or DMC1 was similar to control spermatocytes in mid-late leptotene, and was even higher in mid-zygotene, maybe indicative of the accumulation of unrepaired HR intermediates (Fig. 6a; Extended Data Fig. 6d). The lower percentage of on-axis RAD51 or DMC1 foci that colocalized with RPA2 in cKO spermatocytes compared with control is likely explained by the excess of DSB-independent DMC1 foci (Fig. 6b; Extended Data Fig. 6e). These findings suggest that the efficiency of RAD51 and DMC1 recruitment at sites of meiotic DSBs is not affected by FIRRM and FIGNL1 absence.

**Figure 6.**
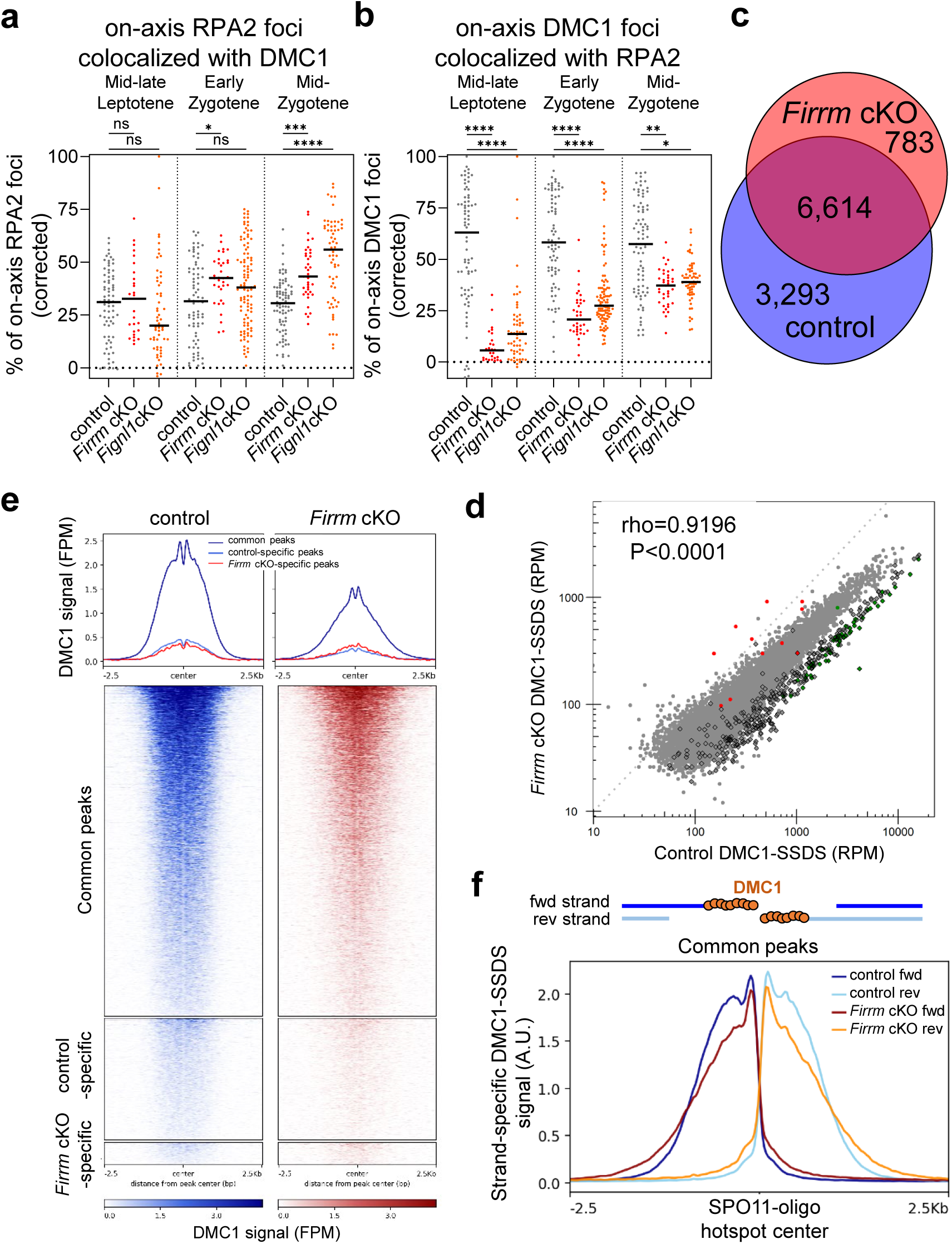
DMC1 is recruited at meiotic DSB hotspots in *Firrm* cKO spermatocytes. **a-b.** Percentages of on-axis RPA2 foci colocalized with on-axis DMC1 foci **(a)**, and vice-versa **(b)** in spreads from mid-late leptotene to mid-zygotene/zygotene-like spermatocyte nuclei from 12-dpp control, *Firrm* cKO and *Fignl1* cKO mice. n=2 mice per genotype. **c.** Numbers of and shared hotspots identified by DMC1-SSDS in spermatocytes from 12 dpp control and *Firrm* cKO mice. **d**. DMC1-SSDS signal correlation between control and *Firrm* cKO mice at hotspots identified in both genotypes. The Spearman rho and associated p-value (two-sided) are shown. Red and green dots indicate hotspots that were significantly over- and under-represented in *Firrm* cKO compared with control spermatocytes (DESeq2, p-value <0.1, log2FC >0 and log2FC <0, respectively). Unchanged autosomal hotspots are represented in gray and chromosome X hotspots by black circled diamonds. **e**. Average plots (top) and corresponding heatmaps (bottom) of DMC1-SSDS intensity (fragments per million, FPM) in control (left) and *Firrm* cKO mice (right) for hotspots that overlap with SPO11-oligo hotspots ^77^ detected in both genotypes (common peaks), in control only (control-specific), or in *Firrm* cKO only (*Firrm* cKO-specific)(see Extended Data Fig. 7a-c). The center of intervals is defined as the center of SPO11-oligo peaks detected in B6 mice, as defined in^77^. **f.** Normalized average distribution of ssDNA type 1 fragments (see Methods) originating from forward (fwd) and reverse strands (rev) at common peaks, defined in **(e)**, for control (dark red, orange) and *Firrm* cKO (dark blue, light blue). The SSDS signal was normalized to have the same cumulated amount of normalized signal over common peaks (on 5-kb windows) for both genotypes.

To assess directly DMC1 recruitment at SPO11-dependent DSB hotspots, we investigated the genome-wide distribution of DMC1-bound ssDNA by chromatin-immunoprecipitation (ChIP), followed by ssDNA enrichment (DMC1-Single Strand DNA Sequencing, SSDS) ^75^ in testes from 12-dpp control and *Firrm* cKO mice. In control mice, the regions enriched in DMC1-bound ssDNA are the ssDNA 3’overhangs that result from DSB resection at meiotic DSB hotspots ^76^. We detected 9,907 peaks in control and 7,397 peaks in *Firrm* cKO spermatocytes (Fig. 6c). Most of these peaks (6,614) were shared. Peaks called specifically in one genotype or the other were most likely shared weakly active hotspots, as inferred from their weak enrichment in both genotypes (Extended Data Fig. 7b). Most of the detected DMC1-SSDS peaks (9,297 out of 10,690) overlapped with previously identified meiotic SPO11-oligonucleotide DSB hotspots (SPO11-oligo hotspots, Extended Data Fig. 7a) ^77^. Moreover, the DMC1-SSDS signal enrichment within peaks was highly correlated in control and *Firrm* cKO samples (Spearman’s rho=0.92; Fig. 6d) with the exception of X chromosome hotspots, which were relatively less enriched in *Firrm* cKO than in control samples (Extended Data Fig. 7d). One possible explanation for this could be a genome-wide accumulation of HR intermediates in *Firrm* cKO that would erase the X chromosome-specific higher DMC1-SSDS enrichment due to delayed DSB repair ^20,77,78^. Overall, this indicates that the recruitment of DMC1 on ssDNA at SPO11-dependent DSB hotspots was efficient, with relative hotspot intensities comparable to wild-type meiosis. The lack of *Firrm* cKO-specific peaks and the efficiency of hotspot detection suggest that DMC1 is not massively associated with ssDNA in non-hotspot regions in *Firrm* cKO testes. The recent finding by Ito and col. that no RAD51-SSDS peaks were detected in *Spo11* KO *Fignl1* cKO testes supports this conclusion ^79^.

We then asked whether FIRRM depletion alters DMC1 extension on resected DSB ends at DSB hotspots. To characterize precisely the DMC1-SSDS signal distribution across DSB hotspots, we defined the center of overlapping SPO11-oligo hotspots as the center of DMC1-SSDS peaks ^77^. This improved significantly the quality of the average DMC1-SSDS signal profile, revealing a non-identical distribution in control and *Firrm* cKO (Fig. 6e; Extended Data Fig. 7c). To facilitate the comparison between control and *Firrm* cKO DMC1-SSDS signal profiles, we normalized the overall signal intensity within common peaks and plotted the resulting normalized strand-specific profiles (Fig. 6f). The distribution of DMC1-SSDS signal in control samples was consistent with previous characterization in wild-type spermatocytes. It is characterized by a spike adjacent to the center of SPO11-oligo hotspots, followed by a shoulder, which is thought to reflect the regulation of the length of resected ssDNA tails covered by DMC1 that measure at least 300 bases and 850-900 bases on average^20,77^. In *Firrm* cKO samples, the DSB-proximal spike was present. However, the density of DMC1-SSDS signal started to decrease right next to the spike, without forming a shoulder. This suggests that the regulation of the length of DMC1-coated ssDNA is altered in the absence of FIGNL1-FIRRM. This alteration of DMC1-SSDS profile was not dependent on hotspot strength (Extended Data Fig. 7e), and was also detected at X chromosome hotspots, suggesting that this is not a consequence of delayed DSB repair (Extended Data Fig. 7d). The presence of the DSB-proximal spike suggests that DMC1 recruitment at DSB sites remains efficient on a short DSB-proximal interval close to the 3’ end of ssDNA tails independently of FIRRM. Conversely, the loss of the shoulder indicates that the mechanism controlling the DMC1 filament length requires FIRRM for full efficiency. One possible scenario is that the FIGNL1-FIRRM complex controls the balance between DMC1 and RAD51 loading on ssDNA. Alternatively, we cannot exclude that the extent of DSB resection is altered.

### *Firrm* cKO is epistatic to *Swsap1* for controlling RAD51 and DMC1 loading

In mouse meiosis, the Shu complex components SWSAP1, SWS1 and SPIDR are required for assembling normal numbers of RAD51 and DMC1 foci, which are 2-to 3-fold fewer in *Swsap1^-/-^*than in wild-type leptotene-zygotene spermatocytes ^36,40–42^. FIGNL1 depletion suppresses the defect of human SWSAP1-depleted cells in forming DNA damage-induced RAD51 foci, suggesting that SWSAP1 antagonizes the anti-RAD51 activity of FIGNL1 ^40^. We generated *Swsap1^-/-^ Firrm* cKO and *Swsap1^-/-^ Fignl1* cKO double mutant mice to determine if the defect in forming meiotic RAD51 and DMC1 foci in *Swsap1^-/-^*spermatocytes depends on FIGNL1-FIRRM. We found that synapsis was defective and meiosis did not progress further than the zygotene-like stage with partial, partly non-homologous synapses in *Swsap1^-/-^ Firrm* cKO and *Swsap1^-/-^ Fignl1* cKO spermatocytes, like in *Firrm* cKO and *Fignl1* cKO single mutants. A small subset of nuclei progressed to pachynema, as observed for *Swsap1^-/-^* spermatocytes, most likely due to incomplete *Firrm* or *Fignl1* deletion. Whereas the number of DMC1 foci was strongly reduced in *Swsap1^-/-^*spermatocytes, double mutant spermatocytes accumulated DMC1, like *Firrm* cKO and *Fignl1* cKO spermatocytes (mean, 123, 30, 247 and 311 DMC1 foci in control, *Swsap1^-/-^*, *Fignl1* cKO and *Swsap1^-/-^ Fignl1* cKO early zygotene spermatocytes, respectively; Fig. 7a-c, Extended Data Fig. 8a-b). Because DMC1 accumulation in *Firrm* cKO and *Fignl1* cKO spermatocytes was essentially DSB-independent (see above), this observation indicates that the formation of DSB-independent DMC1 foci does not require SWSAP1 in the absence of FIGNL1-FIRRM.

**Figure 7.**
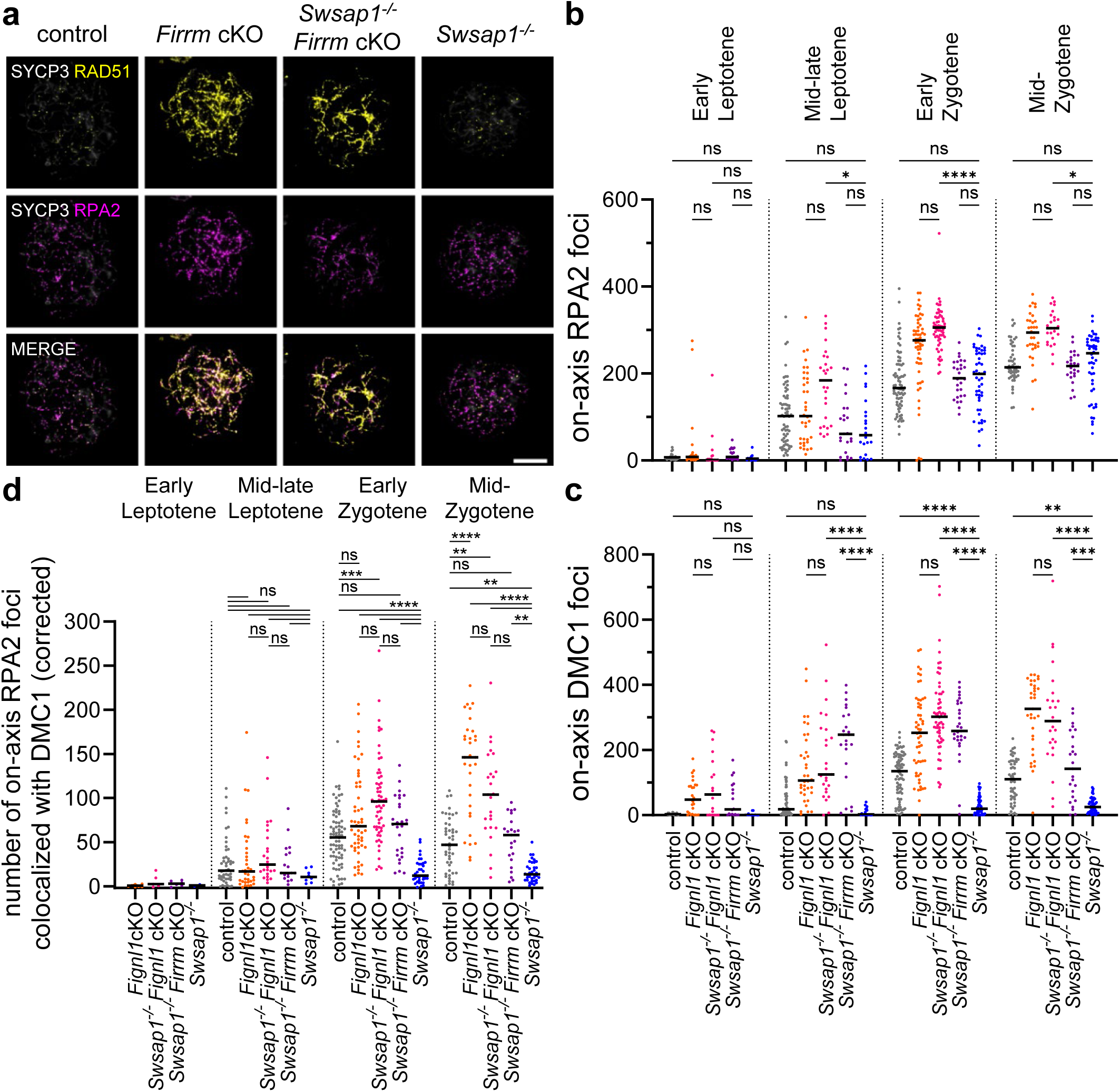
*Firrm* and *Fignl1* deletions restore DMC1 loading in *Swsap1^-/-^* spermatocytes. **a.** Spreads of control, *Firrm* cKO, *Swsap1^-/-^ Firrm* cKO and *Swsap1^-/-^* early zygotene spermatocytes stained for SYCP3 (gray), RAD51 (yellow) and RPA2 (magenta). Scale bar, 10 µm. **b-d.** Total numbers of on-axis RPA2 **(b)** or DMC1 **(c)** foci, or number of on-axis RPA2 foci colocalized with on-axis DMC1 foci (corrected for the numbers expected by chance) **(d)** in spreads from control (n=3 mice), *Fignl1* cKO (n=1), *Swsap1^-/-^ Fignl1* cKO (n=1), *Swsap1^-/-^ Firrm* cKO (n=1) and *Swsap1^-./-^* (n=2) spermatocytes from 17 dpp mice.

We then asked specifically whether SWSAP1 is required for recruiting DMC1 at meiotic DSB sites in the absence of FIRRM or FIGNL1, by examining on-axis colocalized DMC1 and RPA2 foci, used as a proxy for a subset of meiotic DSB sites as described above (Fig. 6a-b; Extended Data Fig. 6a-b). In *Swsap1^-/-^* spermatocytes, only a small fraction of RPA2 foci colocalized with DMC1, in keeping with the reduced number of detectable DMC1 foci. In contrast, the number of RPA2 foci that colocalized with a DMC1 focus was as high in *Swsap1^-/-^ Firrm* cKO and *Swsap1^-/-^ Fignl1* cKO double mutants as in *Firrm* cKO and *Fignl1* cKO single mutants and in control spermatocytes (Fig. 6a, 7d; Extended Data Fig. 8c-e). This suggests that the formation of a normal number of detectable DSB-associated DMC1 nucleofilaments does not require SWSAP1 when FIGNL1 and FIRRM are absent, and is consistent with the hypothesis that SWSAP1 and FIGNL1-FIRRM have antagonistic roles in regulating the formation of stable RAD51/DMC1 nucleofilaments. However, it remains unknown whether these nucleofilaments, which are assembled without counterbalanced regulations by the SWSAP1-containing complex and FIGNL1-FIRRM, are functional for HR.

### FIGNL1 perturbs the architecture of RAD51/DMC1 nucleoprotein filaments and inhibits RAD51- and DMC1-mediated D-loop formation *in vitro*

To determine the HR step(s) in which FIGNL1-FIRRM might be involved, we examined *in vitro* the effect of adding FIGNL1 on the assembly and stability of RAD51 and DMC1 nucleofilaments, and on their subsequent strand invasion activity. We incubated pre-formed RAD51 or DMC1 filaments assembled on a 400 nucleotide (nt) ssDNA or a 400 bp dsDNA with purified human FIGNL1ΔN, in which N-terminal 284 aa were deleted^40^ (Extended Data Fig. 1b, 9a). FIGNL1ΔN did not promote RAD51 and DMC1 displacement from DNA (electrophoretic mobility shift assay in Fig. 8a-b, pre-formed nucleofilament), but induced the formation of a higher molecular weight complex, suggesting that FIGNL1ΔN binds to RAD51/DMC1-DNA filaments. When we mixed FIGNL1ΔN with RAD51 or DMC1 before addition to the DNA substrate, we observed a slight increase in the fraction of free dsDNA (but not ssDNA) that was not complexed with RAD51 or DMC1 (Fig. 8a-c, no pre-formed nucleofilament). Whereas this increase was not significant, it might suggest that the presence of FIGNL1ΔN restricts RAD51 and DMC1 binding to DNA and the subsequent filament elongation. We then used transmission electron microscopy (TEM) to analyze the effect on RAD51 filament formation and architecture upon addition of FIGNL1ΔN at same time as RAD51 to a 400 nt ssDNA (Fig. 8d-e). Addition of FIGNL1ΔN induced the formation of super-complexes that contained several bridged or interwoven filaments. Simultaneously, we observed that individual RAD51 filaments not included in the super-complexes were significantly shorter than RAD51 filaments in controls (mean length of 135 versus 175 nm, respectively; Fig. 8f, Extended Data Fig. 9b). We also detected the formation of some very long filaments (more than 450 nm and up to 3-4 µm). Their length was not compatible with the length of the used DNA substrate, suggesting a DNA-independent polymerization in the presence of FIGNL1ΔN. Consistent with this hypothesis, similar structures were detected by incubating RAD51 with FIGNL1ΔN without adding any DNA (Extended Data Fig. 9b-c). Similarly, the mean length of RAD51 filaments assembled on a 400 bp dsDNA decreased from 194 nm in control to 137 nm in the presence of FIGNL1ΔN (Fig. 8f). The architecture of DMC1 filament assembled both on ssDNA and on dsDNA displayed qualitatively similar alteration (Extended Data Fig. 9d). Altogether, these results show that FIGNL1ΔN limits RAD51/DMC1 assembly on ssDNA and also dsDNA, and affect the filament architecture. We then tested whether these filaments could pair with homologous donor dsDNA (pUC19 plasmid) in a D-loop assay. Pre-formed RAD51, DMC1, and mixed RAD51-DMC1 filaments mediated the formation of 34, 27 and 22% of D-loop products, respectively. Addition of FIGNL1ΔN during filament assembly led to a decrease in the D-loop yield (Fig. 8g-h). When we titrated FIGNL1ΔN in the D-loop reaction, the yield decreased linearly and significantly (Fig. 8h). This showed that the contacting and pairing with homologous DNA of filaments assembled in the presence of FIGNL1ΔN might be affected. This indicates that by limiting the assembly of RAD51 and/or DMC1 on DNA, FIGNL1 could negatively regulate the next strand invasion step required for HR.

**Figure 8.**
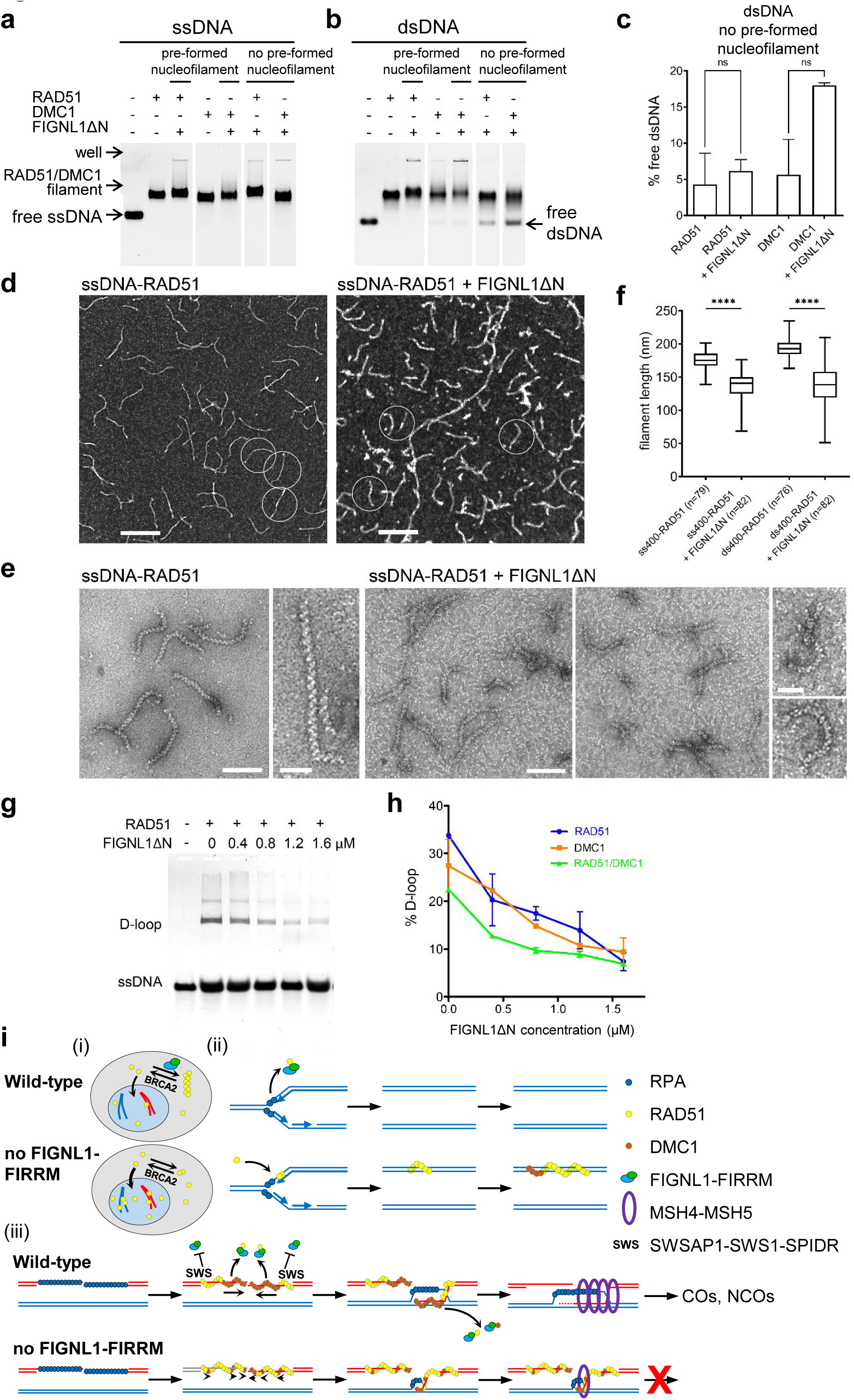
FIGLN1 alters the architecture and the activity of RAD51 and DMC1 nucleoprotein filaments. **a-b.** Electrophoretic Mobility Shift Assay (EMSA). 1 µM RAD51 or DMC1 was incubated (20 minutes) with 3 µM (nucleotide concentration) of a Cy5-labeled 400 nt ssDNA fragment **(a)** or a Cy5-labeled 200 bp dsDNA fragment **(b)** with or without 1.6 µM human FIGNL1ΔN. For the pre-formed nucleofilament panels, RAD51 or DMC1 was incubated with DNA for 5 minutes before adding FIGNL1ΔN for 15 minutes. For the no pre-formed filament panels, RAD51 or DMC1 was added to the reaction concomitantly with FIGNL1ΔN. **c.** Quantification of free dsDNA in the EMSA performed with dsDNA and without pre-formed nucleofilament shown in **(b).** n=2 per condition. Paired t-test, two-sided. **d-f**. Representative TEM images in positive **(d)** and negative staining **(e)** and length distribution **(f)** of RAD51 filaments assembled on 400 nt ssDNA fragments (ss400) without (left, ss400-RAD51) or with human FIGNL1ΔN (right, ss400-RAD51 + FIGNL1ΔN). Some very long filaments (>450nm) that formed in the presence of FIGNL1ΔN **(d)** were not included in the quantification in **(f)** (see Extended Data Fig. 10b). **g-h**. FIGNL1ΔN inhibits the formation of a D-loop by RAD51 and DMC1 *in vitro*. Representative gel (RAD51 in the presence of increasing concentrations of FIGNL1ΔN, from 0.4 to 1.6 µM) **(g)**. Titration of FIGNL1ΔN **(h)** in the D-loop assay. **i.** Model for possible (and non-exclusive) roles of the FIGNL1-FIRRM complex in regulating RAD51 and DMC1 in mouse spermatocytes. (i) The FIGNL1-FIRRM complex may limit the nuclear RAD51 level by sequestering a cytoplasmic RAD51 pool, possibly by promoting RAD51 polymerization, thus preventing its mobilization by BRCA2. (ii) The FIGNL1-FIRRM complex might prevent the stabilization of transient dsDNA-RAD51 association at replication forks during premeiotic replication. (iii) During meiotic recombination, the FIGNL1-FIRRM complex might first promote indirectly the polymerization of a continuous DMC1 filament on the meiotic DSB 3’ ssDNA overhang by preventing the loading of stable RAD51 patches on the 3’ region of the ssDNA tails. This would support the 5’ to 3’ polymerization of DMC1 (arrows) up to the 3’ ends. A factor (e.g., the SWSAP1-SWS1-SPIDR complex) may protect the RAD51 filament from FIGNL1-FIRRM-dependent dissociation in the dsDNA-proximal region of ssDNA tails. The formation of shorter/patchy DMC1 filaments in the absence of the FIGL1-FIRRM complex might not be fully functional for homology search, strand invasion and D-loop stabilization. Post-strand invasion, the FIGNL1-FLIP complex might also be involved in removing RAD51/DMC1 from invading ends involved in intersister (not shown) and/or interhomolog interactions.

## Discussion

The AAA-ATPase FIGNL1 (which plant ortholog is FIGL1) and its partner FIRRM (FLIP or MEICA in plants) were identified recently as negative regulators of meiotic COs in plants ^44,52–54^, and as negative regulators of RAD51 in human cells ^40,43,47–50,57,80^. However, their role in mammalian meiosis remained unknown, although a role for *Fignl1* was hypothesized in mouse male meiosis ^81^. Here, by characterizing male germ line-specific *Fignl1* and *Firrm* cKO mouse models, we uncovered two roles of the FIGNL1-FIRRM complex in male meiosis. First, FIGNL1 and FIRRM are required for meiotic DSB repair and for homologous chromosome synapsis during meiotic prophase I, and thus are essential for male mouse meiosis. Second, the FIGNL1-FIRRM complex prevents DNA damage-independent accumulation of RAD51 and DMC1 on chromatin and chromosome axes in spermatocyte nuclei during premeiotic replication and meiotic prophase I. Similar conclusions were reached in recent studies by characterizing the *Fignl1* cKO ^79^ and the *Firrm* cKO ^57^ mice.

Our data show that FIGNL1 and FIRRM act as negative regulators of RAD51 and DMC1 during meiotic recombination, a function evolutionarily conserved from plants to mammals. However, the role of FIGNL1-FIRRM is much more crucial in mouse spermatogenesis than in *A. thaliana* and rice meiosis where homologous chromosome synapsis and formation of ZMM-dependent type I COs are almost normal in FIGNL1 and FIRRM mutants ^44,52–55^. Plants and mammals show differences in their requirement of specific HR pathways for meiotic DSB repair, homologous chromosome synapsis and progression through meiotic cell cycle. These processes require DMC1, MSH4 and MSH5 in the mouse ^12–14,67,82^. Conversely, in *A. thaliana* and rice, meiotic DSBs are repaired by RAD51-dependent intersister HR in *dmc1* mutants, homologous chromosome synapsis does not depend on MSH4-MSH5, and *dmc1*, *msh4* and *msh5* mutant cells progress through meiotic prophase (reviewed in ^83^). The difference between mice and plants in FIGNL1-FIRRM requirement might be functionally related to these other differences. However, mouse *Fignl1* cKO and *Firrm* cKO spermatocytes also displayed defects not seen in plants, especially a massive, DNA damage-independent RAD51 and DMC1 accumulation and defects in MSH4 focus formation. This suggests that the FIGNL1-FIRRM complex has additional functions in the mouse within the shared framework of RAD51 and DMC1 negative regulation.

We found that in *Fignl1* cKO and *Firrm* cKO spermatocytes, MSH4 focus formation and meiotic DSB repair were impaired, RAD51 and DMC1 foci accumulated at unrepaired DSB sites, and homologous synapsis was defective. These defects have been described in mutants in which strand invasion is impaired (e.g. *Dmc1^-/-^* mice that accumulate only RAD51, *Hop2^-/-^*, *Mnd1^-/-^* mice)^5,6,67,82,84–86^ and in mutants in which strand invasion might be preserved but the HR intermediates are not efficiently stabilized (e.g. *Hrob^-/-^*, *Mcm8^-/-^*, *Mcmd2^-/-^*, *Msh4^-/-^*, *Msh5^-/-^* mice) ^12–14,87–90^. By altering the stability or architecture of the nucleoprotein filament formed by RAD51/DMC1 on ssDNA and/or dsDNA, the FIGNL1-FIRRM complex might play roles before and/or after strand invasion. In the case of a post-strand invasion role, this complex might favor RAD51/DMC1 dissociation from dsDNA in the D-loop, a step required for initiating DNA synthesis to extend the invading strand. In *S. cerevisiae*, the motor protein Rad54 and its paralog Rdh54 are involved in removing RAD51/DMC1 from dsDNA following D-loop formation ^28,32^. In the mouse, the meiotic function of RAD54 and its paralog RAD54B is not crucial because *Rad54 Rad54b* double mutant mice are fertile, although they display persistent RAD51 foci during meiotic prophase ^33,34^. Thus, the FIGNL1-FIRRM complex might be one additional factor that can disassemble RAD51 and DMC1 from dsDNA after D-loop formation. In our *in vitro* assay, human FIGNL1ΔN could not dissociate pre-formed RAD51/DMC1 filaments; however, the full-length FIGNL1-FIRRM complex might possess a stronger activity sufficient to dissociate RAD51/DMC1 efficiently. Alternatively, FIGNL1-FIRRM complex-dependent RAD51/DMC1 filament alteration might render it sensitive to dismantling by other factors. In addition to normal HR intermediate processing, the FIGNL1-FIRRM complex might also dissociate unproductive or potentially toxic post-synaptic RAD51/DMC1 filaments, such as multiple strand invasion or invasion on non-allelic repeated sequences ^91,92^.

In *Firrm* cKO spermatocytes, the average DMC1-SSDS signal profile at meiotic DSB hotspots was altered in a way that suggests that FIRRM is involved in regulating the length of DMC1-ssDNA filaments before strand invasion. In wild-type mouse spermatocytes, the profile of DMC1-SSDS coverage at DSB hotspots and super-resolution microscopy observations indicate that DMC1 typically occupies the DSB-proximal two-third of the DSB 3’ ssDNA end, and RAD51 the DSB-distal third of the same DSB 3’ ssDNA end ^5,6,16,20,73,77^. Because interhomolog recombination is thought to rely solely on DMC1 catalytic activity during meiosis ^18–20^, defects in regulating the length or the continuity of the active DMC1 filament may affect the efficiency of interhomolog search, the formation of a D-loop that can be stabilized by MSH4-MSH5, and homologous chromosome synapsis. Several non-exclusive hypotheses can explain how the FIGNL1-FIRRM complex regulates the DMC1 filament on DSB 3’ ssDNA tails. First, RAD51 nuclear fraction was increased in *Fignl1* cKO and *Firrm* cKO testes (Fig. 1c; Extended Data Fig. 2e), thus RAD51 might outcompete DMC1 on ssDNA tails in these mutants. Mechanistically, a balance between FIGNL1-FIRRM and BRCA2 might regulate the retention of RAD51 in the cytoplasm and contribute to fine-tune RAD51 nuclear level (Fig. 8i, (i)). Indeed, we found that purified human FIGNL1ΔN favors the formation of DNA-independent RAD51 filaments *in vitro* (Extended Data Fig. 9c), while it has been proposed that BRCA2 promotes RAD51 nuclear import by limiting the formation of cytoplasmic RAD51 polymers, which cannot be mobilized ^93^. However, the FIGNL1-FIRRM complex might also play a more direct role in controlling the formation of RAD51 and DMC1 filaments at DSB ssDNA overhangs, based on a model according to which DMC1 filament is seeded at the 3’ end of a RAD51 filament (Fig. 8i, (iii)) ^9^. *In vitro*, RAD51 nucleates randomly on ssDNA tracts, whereas DMC1 prefers to seed at a ds/ssDNA junctions or on a RAD51 patch, and polymerizes in the 5’ to 3’ direction^94^. We thus hypothesize that the FIGNL1-FIRRM complex may disassemble nascent RAD51-ssDNA patches that would otherwise hamper DMC1 filament extension toward the 3’ end of ssDNA tails. In the absence of FIGNL1-FIRRM, the presence of RAD51 patches is predicted to reduce DMC1 occupancy.

Accessory factors might protect RAD51 from the FIGNL1-FIRRM complex on the DSB-distal part of ssDNA tails, permitting preferential RAD51 occupancy specifically in these intervals. One candidate for protecting RAD51 from FIGNL1 is the SWSAP1-SWS1-SPIDR complex ^36,40–42^. Indeed, FIGNL1 interacts with SWSAP1 and SPIDR ^37,40,43^, and SWSAP1 protects RAD51 filaments from FIGNL1 *in vivo* and *in vitro* ^40^. Our observation that SWSAP1 is not required for forming DMC1-RPA2 co-foci in *Swsap1^-/-^ Fignl1*/*Firrm* cKO (Fig. 7d) is consistent with the idea that SWSAP1 supports RAD51/DMC1 focus formation by protecting the nucleofilament from FIGNL1-FIRRM. Interestingly, it was recently reported that in human cells, SWSAP1-SWS1 interacts with the cohesin regulatory protein PDS5B, which localizes to chromosome axes during meiotic prophase ^37,95^. As generally RAD51 localizes closer to the chromosome axis than DMC1 in mouse meiotic prophase (^20,73^ and Fig. 4a), this interaction, if present in meiotic prophase, might provide an anchor that favors preferential RAD51 protection on the DSB-distal part of DSB ssDNA tails. Alternatively, we cannot exclude that DMC1-SSDS profile alteration results from accumulating HR intermediates with a biased DMC1-ssDNA distribution. For example, longer DMC1 filaments might be more frequently engaged in strand invasion, therefore bound on dsDNA and undetectable by ChIP-SSDS, compared with shorter filaments.

In *Fignl1* cKO and *Firrm* cKO spermatocytes, we observed meiotic DSB-independent accumulation of RAD51 foci on chromatin during premeiotic replication that persisted and was accompanied by DMC1 accumulation during meiotic prophase. DNA damage-independent RAD51 foci accumulate in human cells upon RAD51 overexpression ^29^, thus higher RAD51 nuclear concentration in the absence of FIRRM or FIGNL1 might contribute to favor DNA damage-independent RAD51 and DMC1 binding on intact chromatin (Fig. 8i, (i)). In addition, RAD51 and DMC1 DNA damage-independent accumulation is observed in budding yeast and human cells after depletion of RAD54 family DNA translocases ^29–31^. By analogy, the FIGNL1-FIRRM complex might prevent the stabilization of normally transient nascent RAD51-dsDNA filaments at replication forks (Fig. 8i, (ii)). This hypothesis is consistent with our finding that purified human FIGNL1ΔN might reduce RAD51 and DMC1 association with dsDNA *in vitro* (Fig. 8b-c), and with a recent study in human cells showing FIGNL1-FIRRM association with ongoing replication forks in unchallenging conditions ^80^.

The linear RAD51/DMC1 staining detected between SYCP3 synapsed axes suggests that RAD51/DMC1 can associate stably with chromosome axis components, in either a DNA-dependent or DNA-independent manner, in the absence of the FIGNL1-FIRRM complex. DSB-independent RAD51 (but not DMC1) staining along unsynapsed chromosome axes has been previously described in late prophase mouse oocytes ^74,96,97^; however, these structures associating RAD51 and DMC1 along synapsed axes in *Fignl1* cKO and *Firrm* cKO spermatocytes are unusual. RAD51 interacts with several components of meiotic chromosomes, including the axis component SYCP3 ^15^, the SC central element component SYCE2 ^98,99^ and the cohesion regulator PDS5A/B ^100,101^ that interacts also with SWSAP1-SWS1 ^37^. Interestingly, it has been observed by super-resolution microscopy that several cohesin subunits and HORMAD1/2 coat the outside of SYCP3 axis cores ^73,102^, a localization resembling that of RAD51/DMC1 staining between synapsed SYCP3-positive axes. RAD51/DMC1 interactions with components of meiotic chromosome axes might facilitate the accurate HR repair of meiotic DSBs (and incidental DNA damages). In this context, a function of the FIGNL1-FIRRM complex might be to prevent the stabilization of these interactions, other than at DNA damage sites.

Meiotic cells must face the challenge of repairing hundreds of programmed DSBs through several HR pathways, while restricting inappropriate repair that may involve similar HR intermediates. In this study, we started deciphering the functions of the conserved FIGNL1-FIRRM complex in mouse meiosis. We showed that the RAD51/DMC1 filament destabilizing activity of FIGNL1 and FIRRM is implicated in regulating meiotic recombination and restricting inappropriate formation of stable RAD51/DMC1 filaments. Interestingly, although FIGNL1 alters RAD51 and DMC1 filament similarly *in vitro*, it is not clear whether FIGNL1 or FIRRM absence affects DMC1 directly or indirectly through RAD51. The elucidation of the several possible functions of the FIGNL1-FIRRM complex during mouse meiosis will need more *in vitro* and *in vivo* analyses of their functional interactions with other RAD51 and DMC1 regulators.

## Methods

### Mice

All mice used in the study were in the C57BL/6J background. *Firrm^fl/+^*mice (allele *BC055324^tm1c(EUCOMM)Hmgu^*, MGI:5692863) were obtained from the International Knockout Mouse Consortium (IKMC). *Fignl1^fl/+^* mice (allele *Fignl1^tm1c(EUCOMM)Hmgu^*) were generated by Phenomin-Institut Clinique de la Souris (ICS) using the plasmid containing the *Fignl1^tm1a(EUCOMM)Hmgu^* allele (MGI:5287847) obtained from Helmholtz Zentrum München GmbH. *Firrm^fl/fl^* mice were mated with mice that express Cre under the control of the CMV promoter (C57BL/6 Tg(CMV-cre)1Cgn) ^103^ to generate *Firrm*-deleted heterozygous mice (*Firrm^+/-^*). *Firrm^+/-^* mice were mated with Tg(Stra8-icre)1Reb/J (*Stra8-Cre^Tg^*) mice ^59^ to generate *Firrm^+/-^;Stra8-Cre^Tg^* mice. This transgene expresses *Cre* specifically in male germ cells, from undifferentiated spermatogonia to preleptotene spermatocytes ^59^. By crossing *Firrm^fl/fl^* mice with *Firrm^+/-^;Stra8-Cre^Tg^*mice, *Firrm^fl/-^;Stra8-Cre^Tg^* (*Firrm* cKO) and *Firrm^fl/+^*, *Firrm^fl/+^ Stra8-Cre^Tg^* or *Firrm^fl/-^* (*Firrm* control) males were obtained. *Fignl1^fl/-^;Stra8-Cre^Tg^* (*Fignl1* cKO) males were generated using the same strategy as for *Firrm* cKO mice. We assessed the efficiency of germ line CRE-driven excision of the *Firrm^fl^* allele by crossing *Firrm*f*^l/+^ Stra8-Cre^Tg^* males with wild-type females. Among 97 pups, 52 were *Firrm^+/+^* and 45 were *Firrm^+/-^*, with no detection of the *Firrm^fl^* allele in any, indicating an 100% efficiency of transmission of the excised allele to 45 pups. The *Spo11^YF/YF^*^74^ and *Swsap1^-/-^* ^42^ mouse lines were described previously. Primers used for genotyping are listed in Supplementary Table 1. All animal experiments were carried out according to the CNRS guidelines.

### Histology

Mouse testes were fixed in Bouin’s solution for periodic acid-Schiff (PAS) staining at room temperature, overnight. Testes were then embedded in paraffin and 3µm-thick slices were cut. PAS-stained sections were scanned using the automated tissue slide-scanning tool of a Hamamatsu NanoZoomer Digital Pathology system.

### Spermatocyte chromosome spreads

Spermatocyte spreads were prepared with the dry down technique ^104^. Briefly, a suspension of testis cells was prepared in PBS, and then incubated in a hypotonic solution for 8 min at room temperature. Cells were centrifuged, resuspended in 66 mM sucrose solution and spread on slides or coverslips (1.5H, high precision) with 1% paraformaldehyde, 0.05% Triton X-100. Slides/coverslips were dried in a humid chamber for 1-2 h, washed in 0.24% Photoflo200 (Kodak), air-dried, and used for immunostaining or stored at -80°C.

### Immunofluorescence staining

Immunostaining was done as described ^105^. After incubation with a milk-based blocking buffer (5% milk, 5% donkey serum in PBS), spermatocyte spreads were incubated with primary antibodies at room temperature overnight, followed by secondary antibodies (37 °C for 1 h). The antibodies used are listed in Supplementary Table 2. Nuclei were stained with 4’−6-diamidino-2-phenylindole (DAPI, 2 μg/ml) in the final washing step.

For immunostaining with the anti-DMC1 antibody, a specific blocking buffer (0.5% BSA, 0.5% powder milk, 0.5% donkey serum in PBS) was used prior to incubation with the primary antibody. Primary antibody incubation was performed in 10% BSA in PBS. Immunostaining of spermatocyte spreads on coverslips for STED microscopy was done with specific secondary antibodies (Supplementary Table 2), and DAPI was omitted.

### Widefield fluorescent imaging

Widefield images were acquired using one of the following microscopes: Zeiss Axioimager Apotome with 100X Plan Apochromat 1.46 oil DIC objective and 1 ANDOR sCMOS ZYLA 4.2 MP monochrome camera (2048 x 2048 pixels, 6.5µm pixel size) or Zeiss Axioimager 100X Plan Apochromat 1.4 NA oil objective and 1 Zeiss CCD Axiocam Mrm 1.4 MP monochrome camera (1388 x 1040 pixels, 6.45µm pixel size).

### Stimulated emission depletion (STED) super-resolution imaging

Super-resolution images were acquired using a STED microscope (Abberior Instruments, Germany) equipped with a PlanSuperApo 100x/1.40 oil immersion objective (Olympus, Japan). For 3-color STED imaging, immunolabeling was performed using one of the following combinations of secondary antibodies: STAR 460L, STAR ORANGE, STAR RED or STAR GREEN, STAR ORANGE, STAR RED (Supplementary Table 2). STAR 460L and STAR 488 were excited at 485nm, STAR ORANGE at 561nm, and STAR RED at 640nm. Excitation was done with a dwell time of 10µs. STED was performed at 595 nm for STAR 488 and at 775nm for all other dyes. Images were collected in line accumulation mode with detection set at 571-625nm for STAR 460L and STAR ORANGE, 500-580nm for STAR GREEN, and 650-750nm for STAR RED.

### Image analysis

For quantification and colocalization analyses, images were deconvolved using Huygens Professional version 22.10 (Scientific Volume Imaging).

All image analyses were performed using Fiji/ImageJ 1.53t ^106^, with the “MeiQuant” set of tools ^107^ available on github (https://github.com/MontpellierRessourcesImagerie/meiosis_bar).

Single nuclei were cropped manually or using an automatic DAPI signal threshold. Nuclei were sorted into meiotic prophase substages following the criteria described below.

Foci were detected using the Find Maxima function. Focus intensity was determined as the intensity of the maximum for every focus within the chosen category (e.g., on-axis RAD51 foci colocalized with an on-axis DMC1 focus). On-axis and off-axis foci were distinguished on the basis of their localization within (or outside) a binary mask. This ROI was drawn using an automatic SYCP3 axis protein staining threshold (SYCP1 staining was used for MSH4 and TEX11 foci). Because there was no SYCP3 staining-defined axis structure at preleptotene stage, all foci were considered as off-axis foci at this stage.

For two-color focus colocalization, the distance of a given channel focus to the closest second color focus was calculated. Foci were considered as colocalized when this distance was below the minimum resolution distance (0.3µm for widefield images), as in ^108^. The level of random colocalization of foci in channel A (foci A) with foci in channel B (foci B) in any given nucleus was estimated by simulating the random localization of the actual number of foci A within the considered area (on-axis or off-axis as defined above, or within the whole nucleus in preleptotene), and by determining the number of random foci A colocalized with actual foci B. The mean number of colocalizations from 100 simulations was taken as the number of foci A colocalized with foci B by chance in the nucleus (n_random_, “random” on figures), and this was repeated for every nucleus. Reciprocally, the level of random colocalization of foci B with foci A resulted from random simulations of foci B localizations.

In every nucleus, the number of colocalized foci A was corrected for random colocalization by considering that (1) the observed number of colocalized foci A (n_obs_) is composed of one subset of biologically meaningful colocalized foci (“truly” colocalized foci A, n_col_) and one subset of foci A colocalized by chance; (2) the ratio n_random_ /n_T_ (where n_random_ is estimated as described above and n_T_ is the total number of foci A in the nucleus) estimates the frequency of foci A colocalizing by chance among the population of foci A not “truly” colocalized, thus the number of foci A colocalized by chance is (n_T_ – n_col_) * n_random_ /n_tot_, by excluding the truly colocalized foci A from random colocalization.(3) Finally, the estimated number of colocalized foci corrected for random colocalization (n_col,_) was obtained from the formula n_col_=(n_obs_-n_random_)/(n_T_-n_random_), where n_tot_ was the total number of foci counted, n_obs_ the observed number of colocalized foci and n_random_ the mean number of colocalization from 100 simulations as described above. The percentage of corrected colocalization estimate was the ratio of the corrected number of colocalized foci n_col_ over the total number of foci in the same nucleus, n_col_ / n_T_.

For γH2AX quantification, nuclei were cropped manually and the integrated intensity of the γH2AX channel in the cropped region was measured.

Prophase spermatocytes were staged using the following criteria, based on SYCP3 staining. Preleptotene nuclei had patchy weak SYCP3 signal throughout the nucleus. Early leptotene nuclei had focus-like well-defined very short stretches of SYCP3 staining. Mid-late leptotene nuclei had short stretches of SYCP3 fragments. Early zygotene nuclei had longer SYCP3 stretches as the chromosome axes continued to elongate. Mid-zygotene nuclei had very long or full SYCP3 axes, but no or relatively few synapses marked by thicker SYCP3 stretches. Late zygotene had full SYCP3 axes with extensive synapsis marked by thicker SCP3 signal.

### DMC1 chromatin immuno-precipitation, followed by single-strand DNA sequencing (DMC1-SSDS)

DMC1 ChIP-SSDS and library preparation were performed as described in ^109^ using a goat anti-DMC1 antibody (0.5 mg/ml; Santa Cruz, reference C-20). Ten testes from 12 dpp *Firrm^fl/+^;Stra8-Cre^Tg^* (control) and from *Firrm^fl/+^;Stra8-Cre^Tg^* (*Firrm* cKO ) mice were used in each biological replicate. Sequencing was performed on a NovaSeq 6000 PE150 platform in paired end mode (2×150bp).

### Detection of DMC1 ChIP-SSDS peaks

Raw reads were processed using the SSDS-DMC1 Nextflow pipeline ^110^, available on github (https://github.com/jajclement/ssdsnextflowpipeline). Briefly, the main steps of the pipeline included raw read quality control and trimming (removal of adapter sequences, low-quality reads and extra bases) and mapping to the UCSC mouse genome assembly build GRCm38/mm10. Single stranded derived fragments were then identified from mapped reads using a previously published method ^75,111^, and peaks were detected in Type-1 fragments (high confidence ssDNA). To control reproducibility and assess replicate consistency, the Irreproducible Discovery Rate (IDR) method ^112^ was used, following the ENCODE procedure (https://github.com/ENCODE-DCC/chip-seq-pipeline2). The “regionPeak” peak type parameter and default p-value thresholds were used. Briefly, this method performs relaxed peak calling for each of the two replicates (truerep), the pooled dataset (poolrep), and pseudo-replicates that are artificially generated by randomly sampling half of the reads twice, for each replicate and the pooled dataset. Both control and *Firrm* cKO datasets passed the IDR statistics criteria for the two scores (well below 2). By default, the pipeline gave the poolrep as primary output, but for this study the truerep peak sets were considered. Lastly, peak centering and strength calculation were computed using a previously published method ^75^.

The list of SPO11-oligo hotspots from B6 mice and the coordinates (genome build GRCm38/mm10) of their center were from ^77^.

The overlaps between intervals was determined with bedtools ^113^ Intersect on the Galaxy France web interface. For determining overlaps between control and *Firrm* cKO peaks, a minimum overlap of 10%, and reciprocally, was required. The overlap between DMC1-SSDS peaks and the center of SPO11-oligo hotspots ^77^ was considered positive if at least 1 bp of the DMC1-hotspot contained the coordinate of the center of one SPO11-oligo hotspot.

Heatmaps and average plot profiles were generated with deeptools (computeMatrix, plotHeatmap and PlotProfile) on Galaxy France server.

### Preparation of mouse testis protein extracts and western blotting

Cytoplasmic and nuclear extracts were prepared from 12 dpp control, *Firrm* cKO and *Fignl1* cKO mice. Testes were homogenized in hypotonic buffer (10 mM Hepes, pH 7.4, 320 mM sucrose, 0.2 mM PMSF, 1x Complete protease inhibitor cocktail, EDTA-free (Roche), 0.07% beta-mercaptoethanol) in a Dounce homogenizer. After centrifugation (1,000xg at 4°C for 10 min), the supernatant was collected and used as cytoplasmic fraction. The pellet was resuspended in half nuclear packed volume of low salt buffer (20mM Tris-HCl pH7.3, 12.5% glycerol, 1.5mM MgCl_2_, 0.2mM EDTA, 20mM KCl, 1x Complete protease inhibitor cocktail, EDTA-free (Roche), 0.07% beta-mercaptoethanol). Then half nuclear packed volume of high salt buffer (same, but 1.2M KCl) was added drop by drop, incubated at 4°C for 30min with agitation and centrifuged (14,000xg at 4°C for 30 min). The supernatant was collected as nuclear fraction. Cytoplasmic and nuclear fractions were analyzed by western blotting with rabbit anti-FIGNL1 (1/500, Proteintech, 17604-1-AP), rabbit anti-FIRRM (1/500, Abcam, ab121774), rabbit anti-beta tubulin (1/3000, Abcam, ab6046) and guinea pig anti-SYCP3 (1/3,000 ^105^) antibodies. HRP-conjugated secondary antibodies were anti-rabbit IgG-HRP (1:5,000; Cell Signaling Technology) and donkey anti-guinea pig IgG-HRP (1/10,000; Jackson Immuno Research, 706-035-148).

#### Protein purification

Human RAD51 was purified by the CiGEX Platform (CEA, Fontenay-aux-Roses) as follows. His-SUMO-RAD51 was expressed in the *E. coli* strain BRL (DE3) pLys. All protein purification steps were carried out at 4°C. Cells from a 3-liter culture that was induced with 0.5 mM isopropyl-1-thio-ß-D-galactopyranoside (IPTG) at 20°C overnight were resuspended in 1x PBS, 350 mM NaCl, 20 mM imidazole, 10% glycerol, 0.5 mg/ml lysozyme, Compete Protease Inhibitor (Roche), 1 mM 4-(2-aminoethyl)benzenesulfonyl fluoride (AEBSF). Cells were lysed by sonication and the insoluble material was removed by centrifugation at 150,000 x g for 1h. The supernatant was incubated with 5 ml of Ni-NTA resin (Qiagen) for 2h. The mixture was poured into an Econo-Column Chromatography Column (BIO-RAD) and beads were washed first with 80 ml W1 buffer (20 mM Tris HCl pH 8, 500 mM NaCl, 20 mM imidazole, 10% glycerol, 0.5% NP40), followed by 80 ml of W2 buffer (20mM Tris HCl pH 8, 100mM NaCl, 20mM imidazole, 10% glycerol, 1 mM DTT). Then, His-SUMO-RAD51 bound to the beads was resuspended in 8ml of W2 buffer and incubated with SUMO protease at a 1/80 ratio (w/w) for 16 h. RAD51 without the His-SUMO tag was then recovered into the flow thru and directly loaded onto a HiTrap heparin column (GE Healthcare). The column was washed with W2 buffer and then a 0.1-1M NaCl gradient was applied. Fractions containing purified RAD51 were concentrated and dialyzed against storage buffer (20mM Tris HCl pH 8, 50mM KCl, 0.5 mM EDTA, 10% glycerol, 1 mM DTT, 0.5 mM AEBSF) and stored at -80°C. Human RPA was purified by the CiGEX Platform (CEA, Fontenay-aux-Roses) as previously described ^114^.

For human FIGNL1 purification, *FIGNL1ΔN* without the region encoding the N-terminal 284 aa, allowing the production of soluble protein^40^, was inserted into the pET15 vector (Novagene), and the protein was overexpressed in *E. coli* BL21(DE3) cells upon addition of 0.2mM IPTG at 37°C for 3h. Cell pellets were resuspended in buffer A (50mM Tris-HCl pH7.4, 500 mM NaCl, 5% glycerol, 5mM MgCl_2_, 5 mM β-mercaptoethanol, 1 mM PMSF, 0.1% NP40, 20 mM imidazole, cOmplete Protease Inhibitor Cocktail), disrupted by French press (6 bar) and cleared by centrifugation following incubation with the benzonase nuclease (Sigma) at 4°C for 30 min. The supernatant was loaded on a 1 ml HisTrap Fast-Flow column (GE healthcare) and equilibrated with buffer A on an ÄKTA pure system. After a washing step, proteins were eluted with buffer A supplemented with 300 mM imidazole. FIGNL1ΔN was further purified by size exclusion chromatography using a HiLoad 16/600 Superdex 200 column (GE Healthcare) in buffer B (50mM Tris-HCl pH7.4, 200 mM NaCl, 10% glycerol, 5mM MgCl_2_, 5 mM β-mercaptoethanol). The peak fractions were concentrated with Amicon Ultra 30K (Millipore) and stored at -80°C.

#### RAD51 and DMC1 filament electromobility shift assay (EMSA)

RAD51 and DMC1 filaments were formed by incubating 3 μM (nucleotide concentration) of 400 nt ssDNA or dsDNA labeled with Cy5 with 1 μM RAD51 (1 protein per 3 nt) or 1.5 μM DMC1 (>1 protein per 3 nt to obtain fully covered DNA) in a buffer containing 10 mM Tris-HCl pH7.5, 50 mM NaCl, 2 mM MgCl_2_, 2 mM CaCl_2_, 2 mM ATP, and 1 mM DTT at 37°C for 20 min. Then, 1.6 μM of FIGLN1ΔN was added to the reaction to test its effect on filament assembly and architecture (pre-formed filament). Control experiments, in absence of FIGLN1ΔN, were realized by adding the same volume of FIGLN1ΔN storage buffer to ensure that the detected effect is not due to the FIGLN1ΔN buffer components. Alternatively, RAD51 or DMC1 was added concomitantly with FIGLN1ΔN to the reaction (no pre-formed filament). Protein-DNA complexes were fixed in 0.01% glutaraldehyde at room temperature for 5 min. Then, the reaction products were analyzed using 1% agarose gel in 0.5x Tris acetate/EDTA at 4 °C. Images were acquired using a Typhoon imager (GE Healthcare Life Science).

#### Tramsmission electron microscopy (TEM) analysis of RAD51 and DMC1 filaments

RAD51 and DMC1 filaments were formed by incubating 7.5 μM (nucleotide concentration) of 400 nt long ssDNA and dsDNA with 2.5 μM RAD51 (1 protein per 3 nt) or 3.5 μM DMC1 in a buffer containing 10 mM Tris-HCl pH7.5, 50 mM NaCl, 2 mM MgCl_2_, 2 mM CaCl_2_, 2 mM ATP and 1 mM DTT at 37°C for 20 min. Then, 1.6 μM of FIGLN1ΔN was added to the reaction at the same time as RAD51/DMC1. Again, control experiments, in absence of FIGLN1ΔN, were realized by adding the same volume of FIGLN1ΔN storage buffer. For filament length analysis, positive staining combined with a TEM dark-field imaging mode were used: 1 μL of the reaction was quickly diluted 20 times in a buffer containing 10 mM Tris-HCl pH 7.5, 50 mM NaCl, 2 mM MgCl_2_, 2 mM Cacl_2_. During one minute, a 5 μL drop of the dilution was deposited on a 600-mesh copper grid previously covered with a thin carbon film and pre-activated by glow-discharge in the presence of amylamine (Sigma-Aldrich, France) ^115,116^. Grids were rinsed and positively stained with aqueous 2 % (w/v) uranyl acetate, dried carefully with a filter paper. To better observe FIGLN1ΔN effect on the filament architecture, samples were also spread using negative staining and observed in bright-field mode. For this, a drop of the reaction was directly deposited on a carboned copper grid pre-activated with glow discharge (plasma).

TEM grids were observed in the annular dark-field mode in zero-loss filtered imaging or in canonical bright-field imaging using a Zeiss 902 transmission electron microscope. Images were captured at a magnification of 85,000× with a Veleta CCD camera and analyzed with the iTEM software (both Olympus Soft Imaging Solution). For quantification, the filament length was measured in at least two independent experiments with a total of at least 75 molecules measured.

#### D-loop *in vitro* assay

RAD51 and DMC1 filaments were formed in the same conditions as for the EMSA analysis. The same incubation conditions and buffer were used to assemble mixed RAD51/DMC1 filaments by incubating 3 μM (nucleotide concentration) of 400 nt ssDNA substrates with 1.25 μM RAD51 plus 0.75 μM DMC1. In the second step, 15 nM in molecules of homologous dsDNA donor (pUC19 plasmid purified on MiniQ ion exchange chromatography column) was introduced in the reaction and in case of DMC1 filaments, 4 mM more CaCl_2_ was added, and then the mixture was incubated at 37°C for 30 min. The reaction was stopped with 0.5 mg/mL proteinase K, 1% SDS, 12.5 mM EDTA at 37°C for 30 min and separated on 1% TAE agarose gels (80 V, for 30 min).

### Statistical analysis and reproducibility

The statistical analyses of cytological observations were done with GraphPad Prism 9. A contingency chi-square test was used to compare stage distributions. The nonparametric Dunn’s multiple comparison test was used to compare focus counts, colocalized focus counts and fractions, and γH2AX intensity among genotypes. The nonparametric Wilcoxon signed-ranks test was used to compare true colocalization versus random colocalization of foci. All tests, sample size, and p values (n.s., not significant, *P < 0.05, **P < 0.01, ***P < 0.001, ****P < 0.0001) are provided in the corresponding legends and/or figures. If not otherwise stated, at least two animals/genotype were analyzed and similar results were obtained.

## Data availability

The DMC1-SSDS raw and processed data for this study have been deposited in the European Nucleotide Archive (ENA) at EMBL-EBI and are available through the project identifier PRJEB62127.

## Acknowledgements

We would like to thank Masaru Ito and Akira Shinohara for sharing unpublished results, Maria Jasin for *Swsap1* mice, Qinghua Shi for the guinea-pig anti-DMC1 antibody. We thank Thomas Robert for critical reading of the manuscript. We thank the following Montpellier Biocampus facilities for their service: Anne Sutter and the animal facility (RAM) for animal care, Manon Leportier for managing our mouse strains, the Réseau d’Histologie Expérimentale de Montpellier (RHEM) for histology. We acknowledge the support of Marie-Pierre Blanchard for help with STED microscopy and the imaging facility MRI, member of the national infrastructure France-BioImaging infrastructure supported by the French National Research Agency (ANR-10-INBS-04, “Investments for the future”). We are grateful to Xavier Veaute from the CiGEX Platform (CEA, Fontenay-aux-Roses), and to the https://www.france-bioinformatique.fr/ and the https://www.france-bioinformatique.fr/fr/cluster for providing computing resources on Galaxy France.

This work was supported by the Agence Nationale de la Recherche ANR FIRE (ANR-17-426 CE12-0015) to FB, PD, JBC and RK; by the Ligue contre le Cancer (Comité départemental de l’Hérault and Comité départemental du Gard) to FB; by the Ligue contre le Cancer (MB/CB/070-22) to RK. AZ was funded by ANR FIRE (ANR-17-426 CE12-0015) and a PhD fellowship FDT202106012805 from the Fondation pour la Recherche Médicale (FRM).

## Author contributions

AZ, PD, RK, RM and FB conceived and designed the experiments. RM identified FIRRM as the mouse homolog of plant FLIP. RTM provided unpublished information. AZ performed most mouse experiments, SB and FB performed some mouse experiments. AZ and FB interpreted and analyzed mouse data with input from BdM. PD, VR, RK performed, analyzed and interpreted biochemical experiments with contribution from JBC. JC developed the method for image analysis. PA and JAJC developed the bioinformatic pipeline for analyzing SSDS data. JAJC and FB performed bioinformatic analysis of SSDS data. AZ, PD, RK and FB wrote the manuscript with input from all authors.

## Competing interests

The authors declare no competing interests.

---

**Supplementary Table 1.**
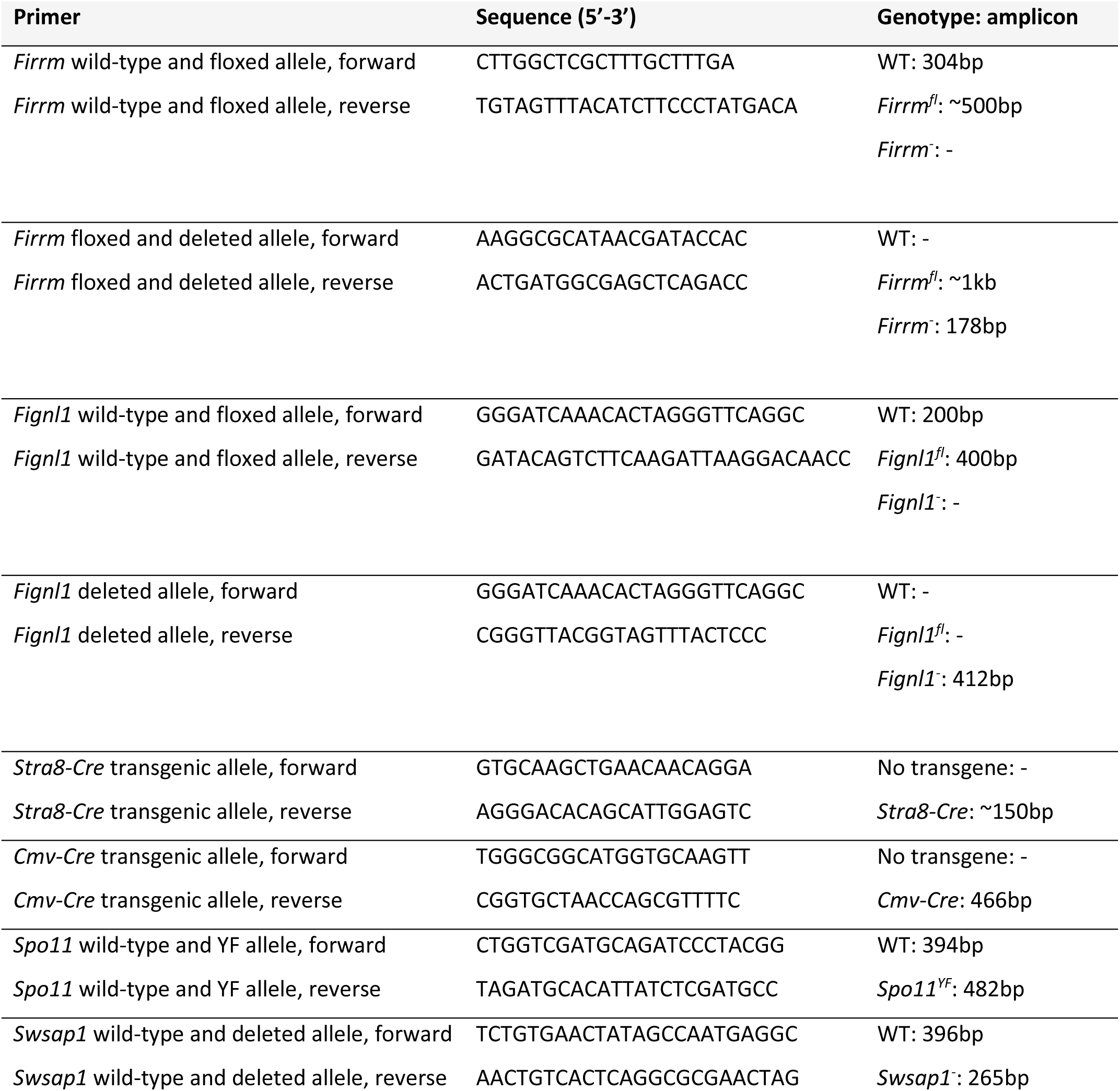
Primers used for mouse genotyping.

**Supplementary Table 2.**
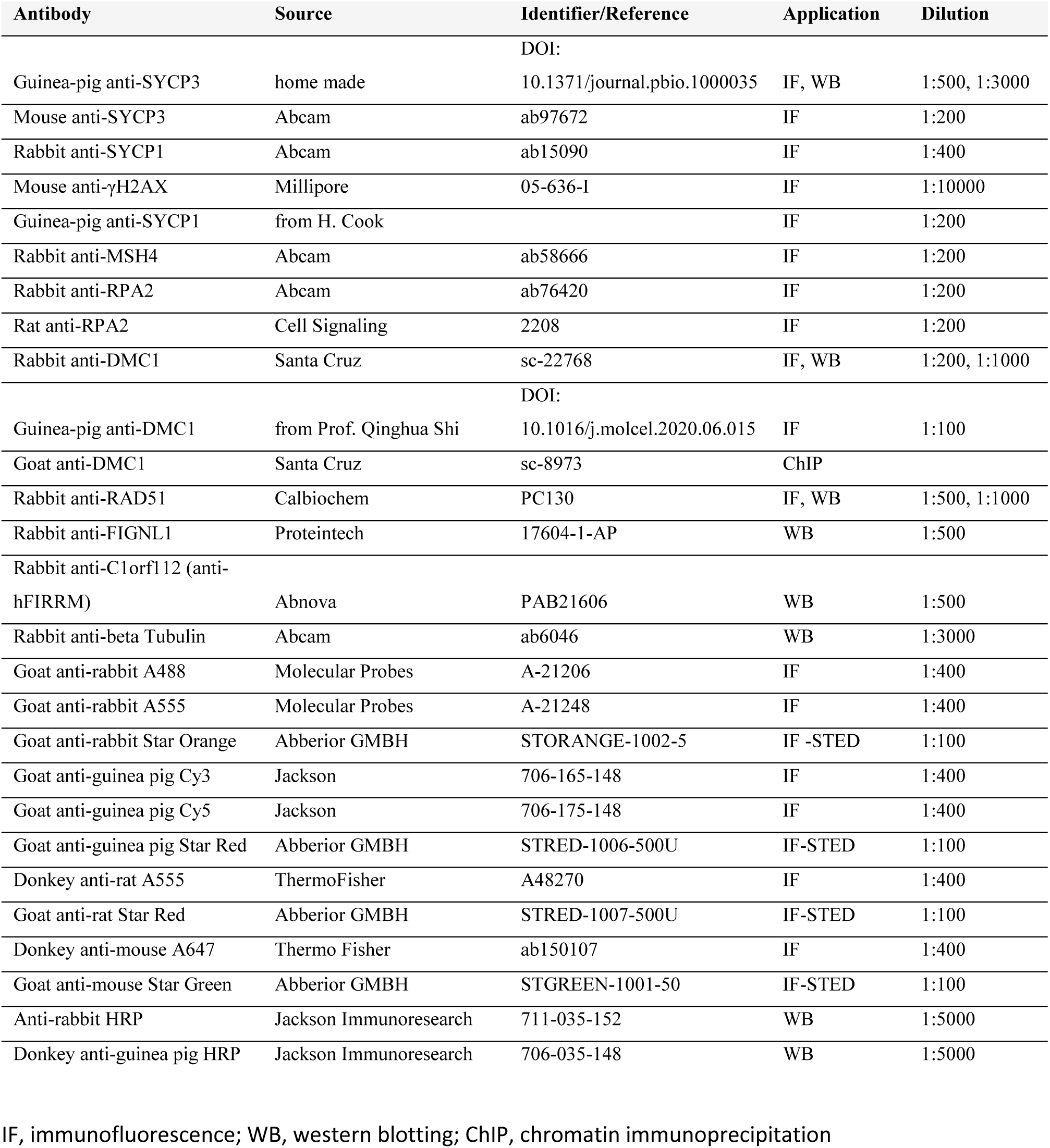
List of antibodies used in this study.

## Figure Legends

**Extended Fig. 1.**
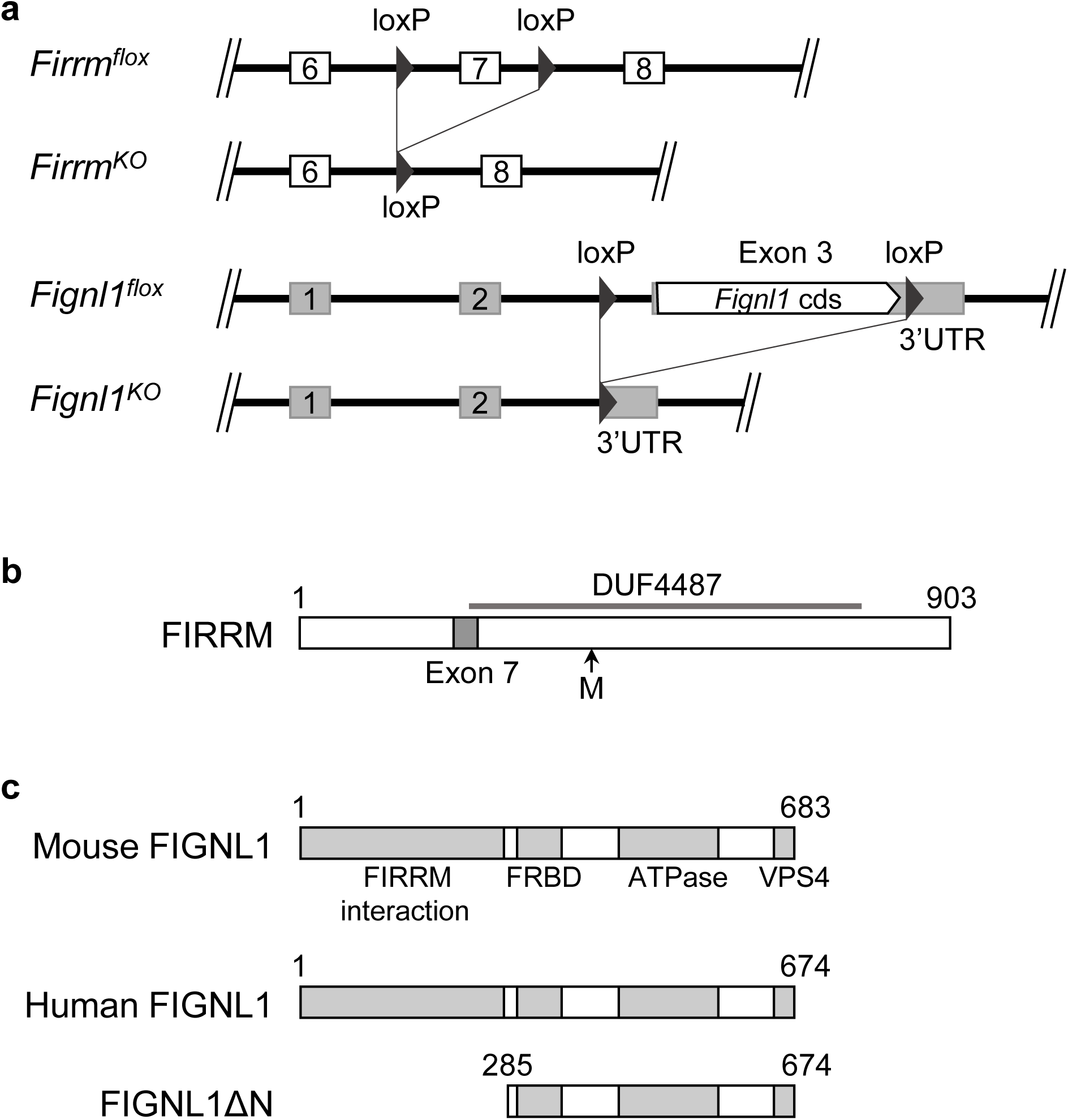
Structure of *Firrm* and *Fignl1* cKO alleles. **a.** Genomic structure of the floxed and knockout (KO) *Firrm* and *Fignl1* alleles. Open boxes, coding exons; gray-filled boxes, non-coding exons**. b.** The mouse FIRRM protein. The conserved DUF4487 domain is indicated, with the position of exon 7 deleted in the KO (generating a frameshift), and the following internal methionine (M, position 406). **c.** Domain organization of mouse (683aa) and human (674aa) FIGNL1 proteins and the human FIGNL1ΔN truncation mutant.

**Extended Fig. 2.**
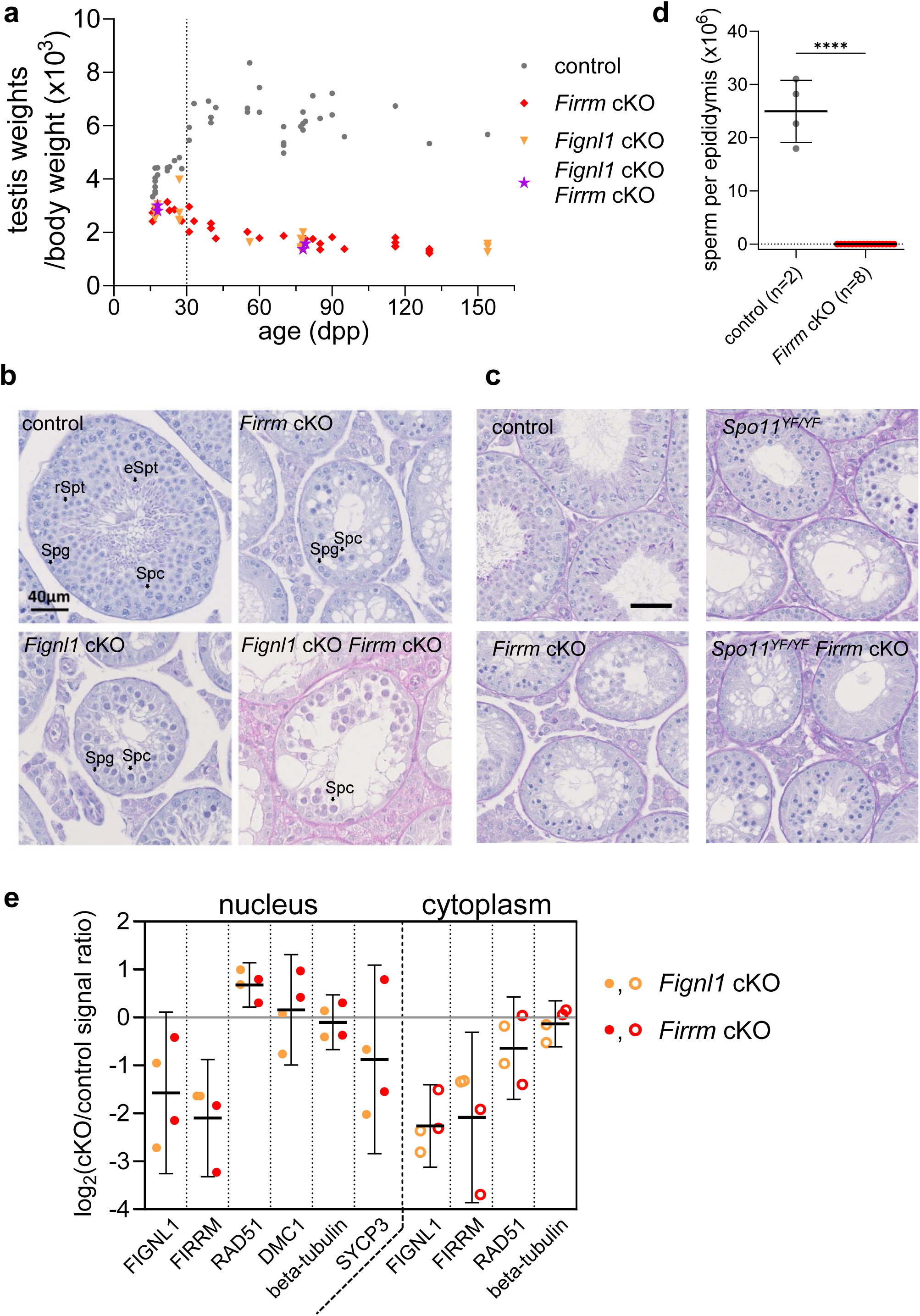
Spermatogenesis is defective in *Firrm* cKO and *Fignl1* cKO mice. **a.** Testis weight relative to body weight in control (n=44), *Firrm* cKO (n=24), *Fignl1* cKO (n=18) and *Firrm* cKO *Fignl1* cKO (n=4) mice of various ages (16 dpp to 154 dpp). Data from 30 dpp (dotted line) or older individuals were plotted in Fig. 1a. **b-c.** Periodic acid-Schiff-stained testis sections from 10-11-week-old **(b)** or 8-week-old **(c)** mice of the indicated genotypes. Note the depletion of germ cells in 10-11-week-old *Firrm* cKO, *Fignl1* cKO and *Firrm* cKO *Fignl1* cKO tubules, and in 8-week-old *Firrm* cKO (and *Spo11^YF/YF^ Firrm* cKO) compared to *Spo11^YF/YF^* tubules. **e.** Sperm count in cauda epididymis from 4-month-old mice (unpaired t-test). **d.** Sperm count in cauda epididymis from 4-month-old mice (unpaired t-test). **e.** Quantification of the western blot signal of cytoplasmic (80µg) and nuclear (100µg) fractions from testes of 12 dpp mice of the indicated genotypes (n=2 independent extracts from 3-4 mice per genotype). The log_2_ of *Fignl1 cKO* or *Firrm cKO* over control signal ratio is shown. Black bar, geometric mean with 95% confidence interval of pooled data from *Fignl1* cKO and *Firrm* cKO experiments.

**Extended Fig. 3.**
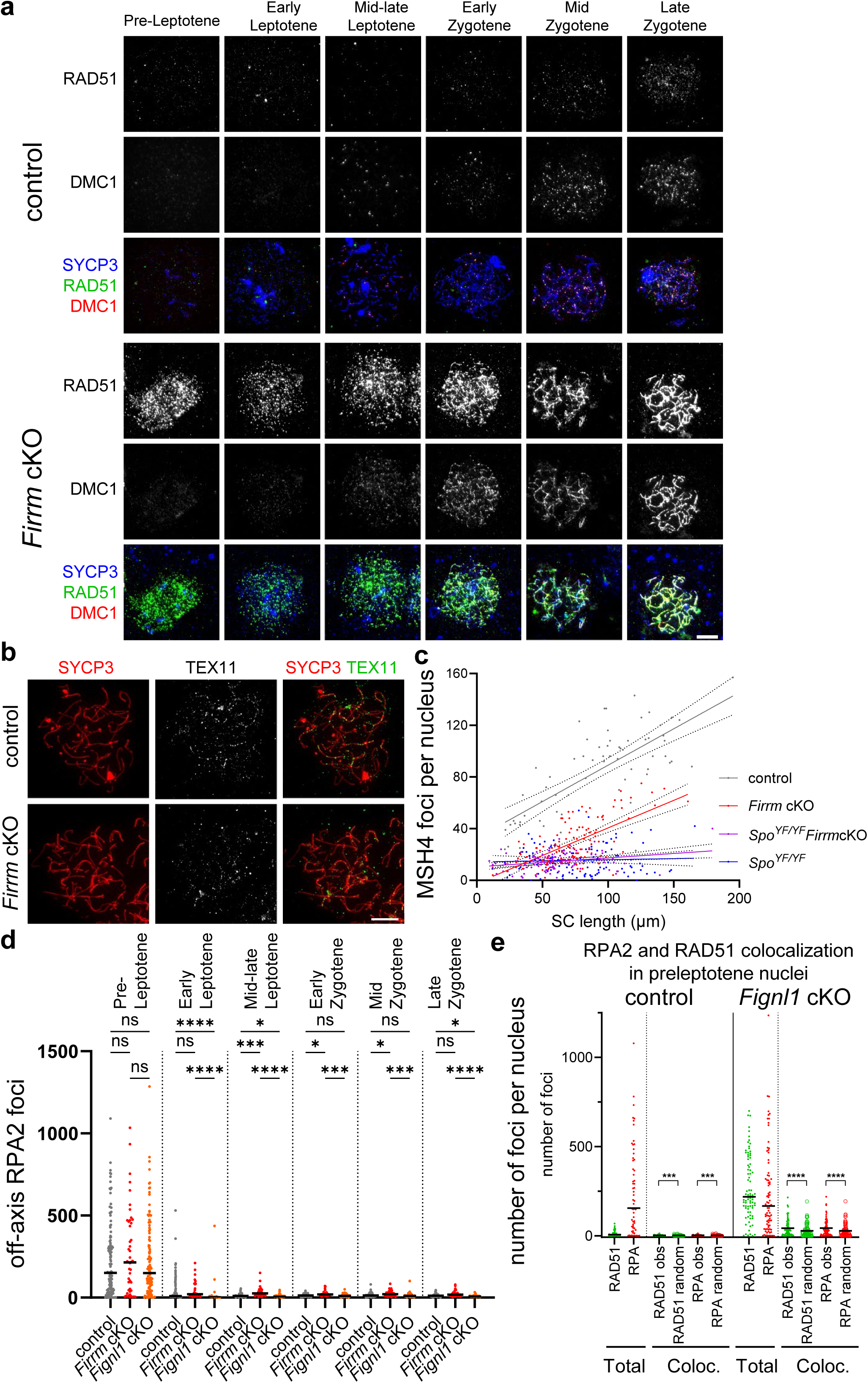
Increased RAD51 and DMC1 loading, and defective MSH4 and TEX11 focus formation, in *Firrm* cKO and *Fignl1* cKO spermatocytes. **a.** Representative images of preleptotene to late zygotene spermatocyte spreads from control and *Firrm* cKO mice stained for SYCP3, RAD51 and DMC1. Scale bar, 10 µm. **b.** Spreads of zygotene spermatocytes from 16 dpp control and *Firrm* cKO mice stained with SYCP3 and TEX11. **c.** Number of MSH4 foci along SYCP1-marked synaptonemal complex fragments in control, *Firrm* cKO, *Spo11^YF/YF^ Firrm* cKO, and *Spo11^YF/YF^*zygotene or zygotene-like spermatocytes. The number of MSH4 foci varied with the SC length in control and *Firrm* cKO spermatocytes. The linear regression fit is shown, with the standard error. **d.** Numbers of off-axis RPA2 foci in control (gray), *Firrm* cKO and *Fignl1* cKO spermatocytes (red). Same mice as in Fig. 2d. **e.** Numbers of all and of colocalized RAD51 (green) and RPA2 (red) foci in spreads of preleptotene control and *Fignl1* cKO spermatocyte nuclei. n=2 (control) or n=3 mice (*Fignl1* cKO). The numbers of observed and expected by chance (random) colocalized foci are shown (see Methods). Wilcoxon two-tailed test.

**Extended Fig. 4.**
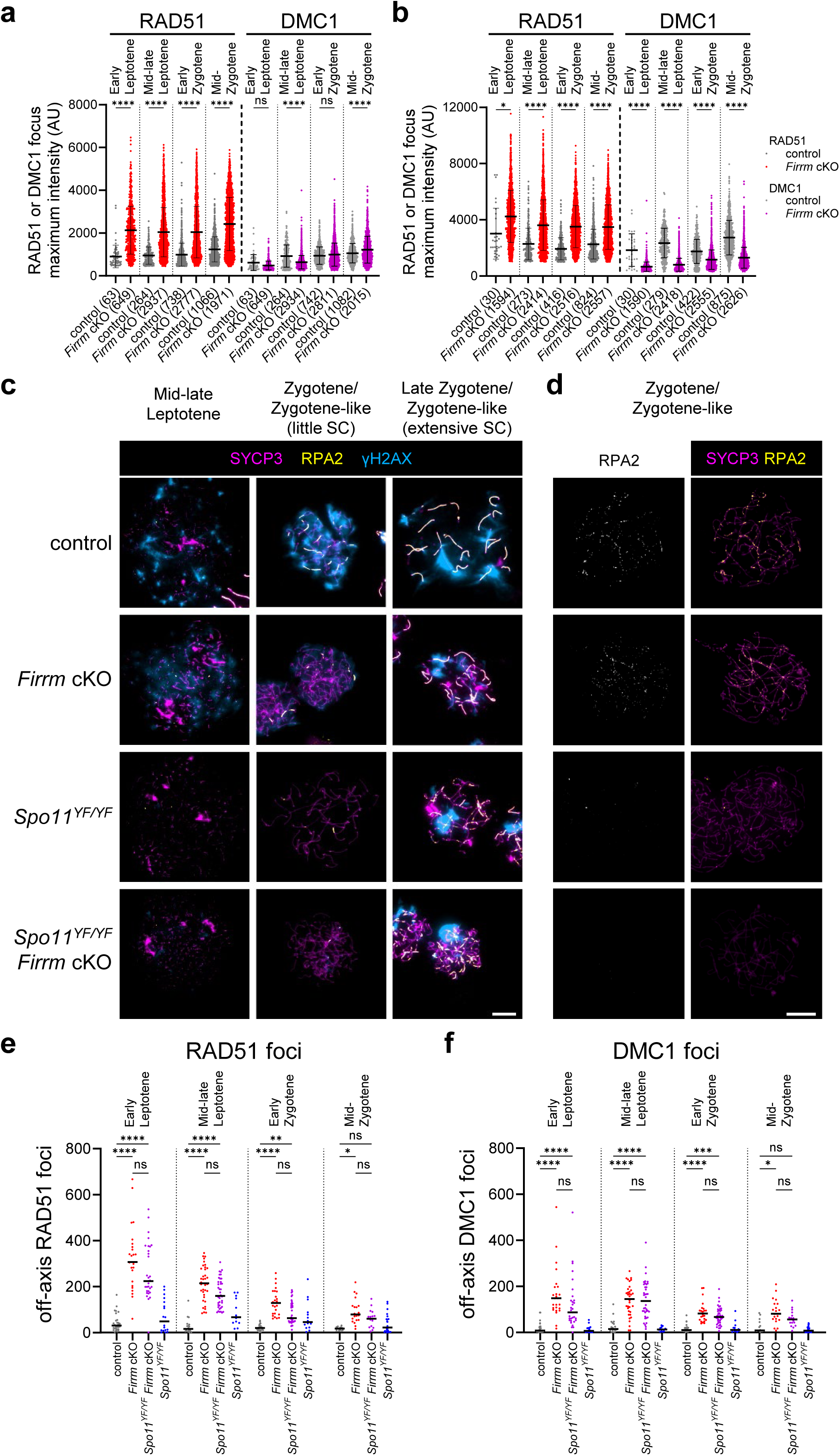
Absence of meiotic DSBs, but presence of RAD51 and DMC1 off-axis foci, in *Spo11^YF/YF^ Firrm* cKO spermatocytes. **a-b.** Intensity of RAD51 and DMC1 foci in spread spermatocytes from 12-dpp mice (replicate 1 **(a)** and replicate 2 **(b)**, n=1 mouse per genotype per replicate) co-stained for RAD51 and DMC1. Only co-localized foci were considered. **c-d.** Representative spread nuclei of spermatocytes from control, *Firrm* cKO, *Spo11^YF/YF^ Firrm* cKO, and *Spo11^YF/YF^* mice stained for SYCP3, SYCP1 and γH2AX **(c)** or for SYCP3 and RPA2 **(d)**. Scale bar, 10 µm. **e-f**. Numbers of off-axis RAD51 **(e)** and DMC1 **(f)** foci for control, *Firrm* cKO, *Spo11^YF/YF^ Firrm* cKO, and *Spo11^YF/YF^* spermatocyte spreads.

**Extended Fig. 5.**
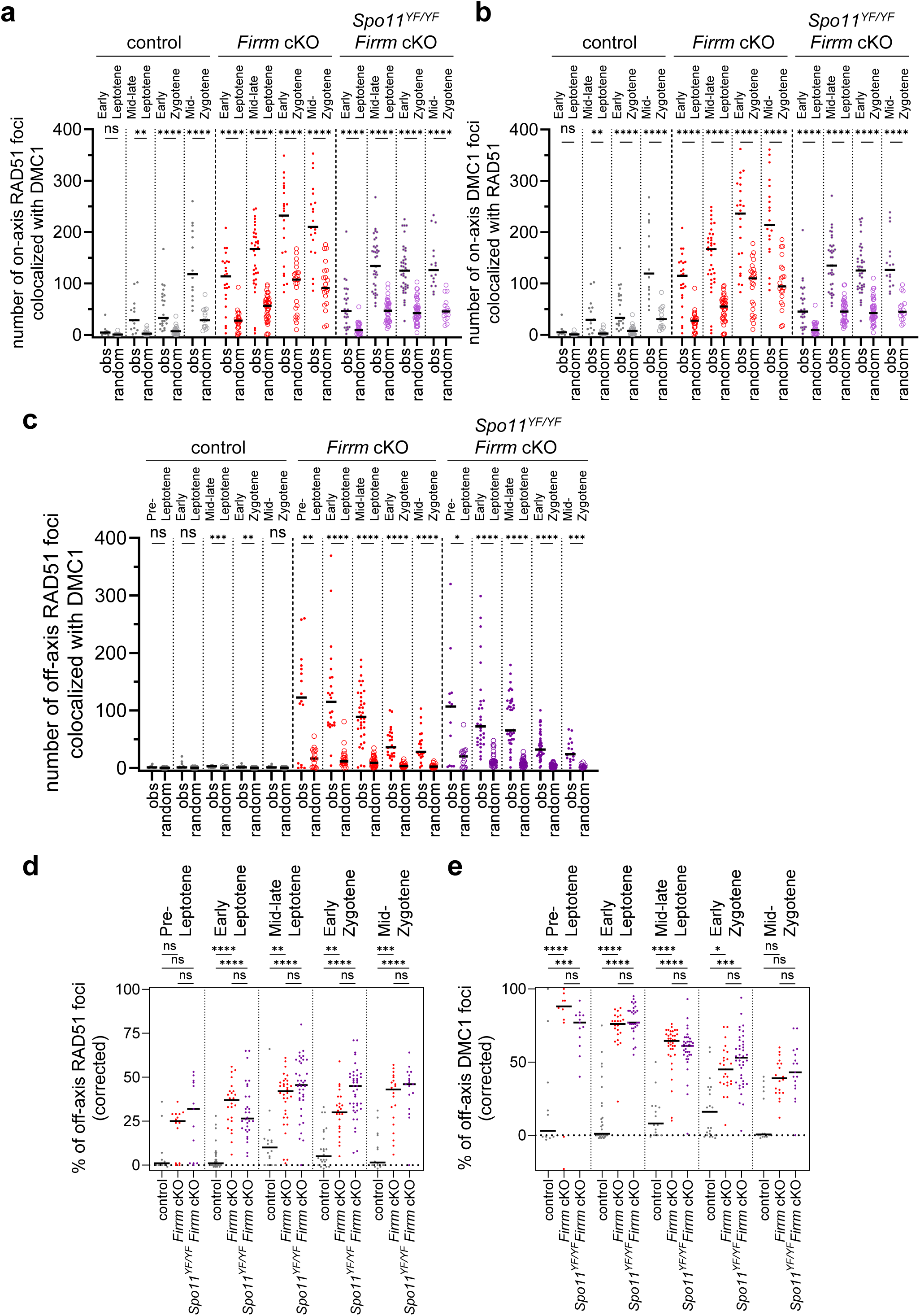
SPO11 DSB-independent DMC1 and RAD51 foci colocalize in *Spo11^YF/YF^ Firrm* cKO spermatocytes. **a-b**. Number of on-axis RAD51 foci colocalized with on-axis DMC1 foci **(a)**, and vice-versa **(b),** on spread from spermatocytes of 12 dpp mice of the indicated genotypes. n=2 mice per genotype **(a-e)**. **c-e**. Number **(c)** or percentage (corrected for random colocalization) **(d-e)** of off-axis RAD51 foci colocalized with off-axis DMC1 foci **(c-d)**, and vice-versa **(e)**.

**Extended Fig. 6.**
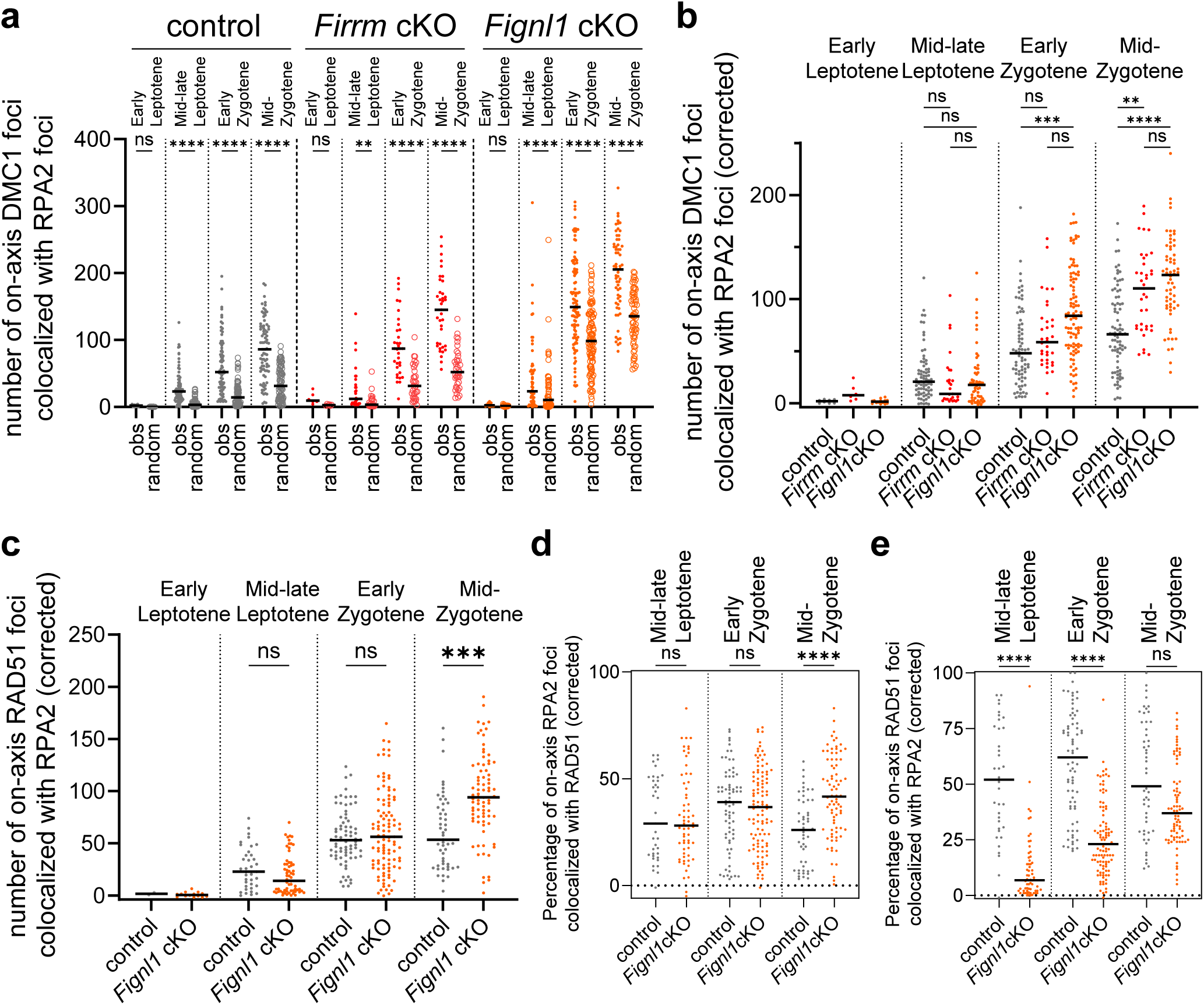
Similar numbers of DMC1 and RPA2 foci colocalize in wild-type and *Firrm* cKO spermatocytes. **a-b.** Number of on-axis DMC1 foci colocalized with on-axis RPA2 foci on spreads from spermatocyte nuclei from control, *Firrm* cKO and *Fignl1* cKO mice. The observed (obs) and expected by chance (random) numbers of DMC1 foci colocalized with RPA2 are shown in **(a)**, while the counts are corrected for the number expected by chance in **(b)**. obs, number of detected colocalizing foci. Random, average of 100 simulations where the colocalization of randomly distributed DMC1 foci with actual RPA2 foci was measured. Wilcoxon two-tailed test **(a)**. **c-e.** Number **(c)** and percentages **(d-e)** of on-axis RAD51 foci colocalized with on-axis RPA2 foci on spreads from spermatocyte nuclei from control (n=2) and *Fignl1* cKO (n=3) mice. The counts corrected for the number expected by chance are shown in **(c)**. **d, e**. Percentage (corrected for random colocalization) of RPA2 foci colocalized with RAD51 **(d)** and vice-versa **(e)**.

**Extended Fig. 7.**
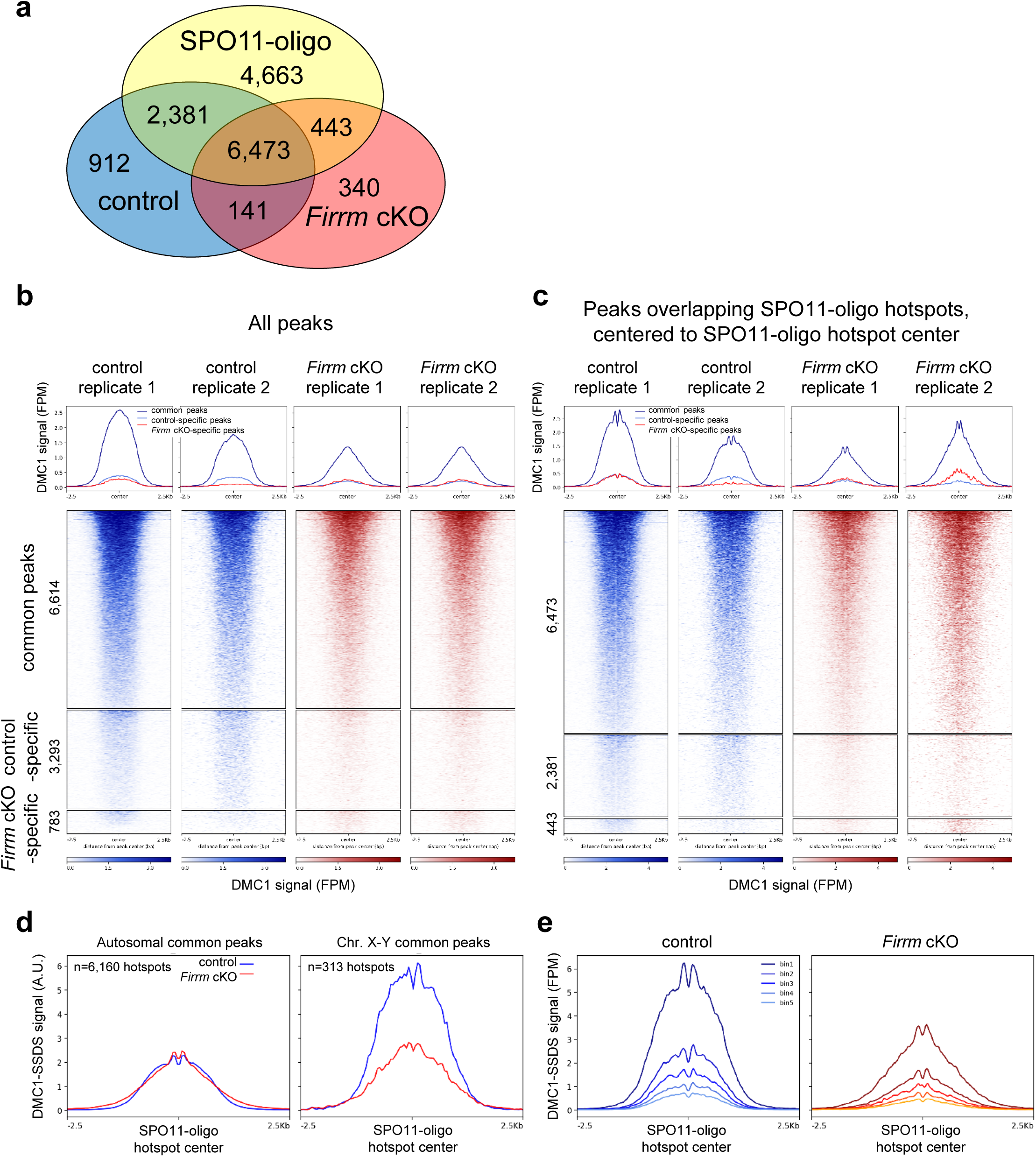
DMC1 recruitment at meiotic DSB hotspots in *Firrm* cKO spermatocytes. **a.** Numbers and overlap of hotspots identified by DMC1-SSDS in spermatocytes from 12 dpp control and *Firrm* cKO mice, and of SPO11-oligo hotspots detected in C57BL/6J mice in ^77^. **b-c**. Average plots (top) and corresponding heatmaps (bottom) of DMC1-SSDS signal in control and *Firrm* cKO mice (2 biological replicates/each), at all common, control-specific, and *Firrm* cKO-specific DMC1 hotspots identified in our analysis **(b)**, and at hotspots overlapping with SPO11-oligo hotspots detected in C57BL/6J mice **(c)**. In **(c)**, the center of the intervals was the center of SPO11-oligo peaks detected in B6 mice, as defined in (Lange, 2016). **d.** Average DMC1-SSDS signal distribution at common DMC1 hotspots, defined in **(c)**, at autosomal hotspots (left panel) and at X and Y chromosome hotspots (right panel), for control (blue) and *Firrm* cKO (red). The DMC1-SSDS signal was normalized to have the same total amount of normalized signal for all common hotspots (on 5-kb windows) in both genotypes. The relative excess of DMC1-SSDS signal at X-Y chromosome hotspots in control is clear. **e.** Average plots of DMC1-SSDS signal intensity (in FPM) at common hotspots defined in **(c)**, ranked within 5 bins of decreasing intensity.

**Extended Fig. 8.**
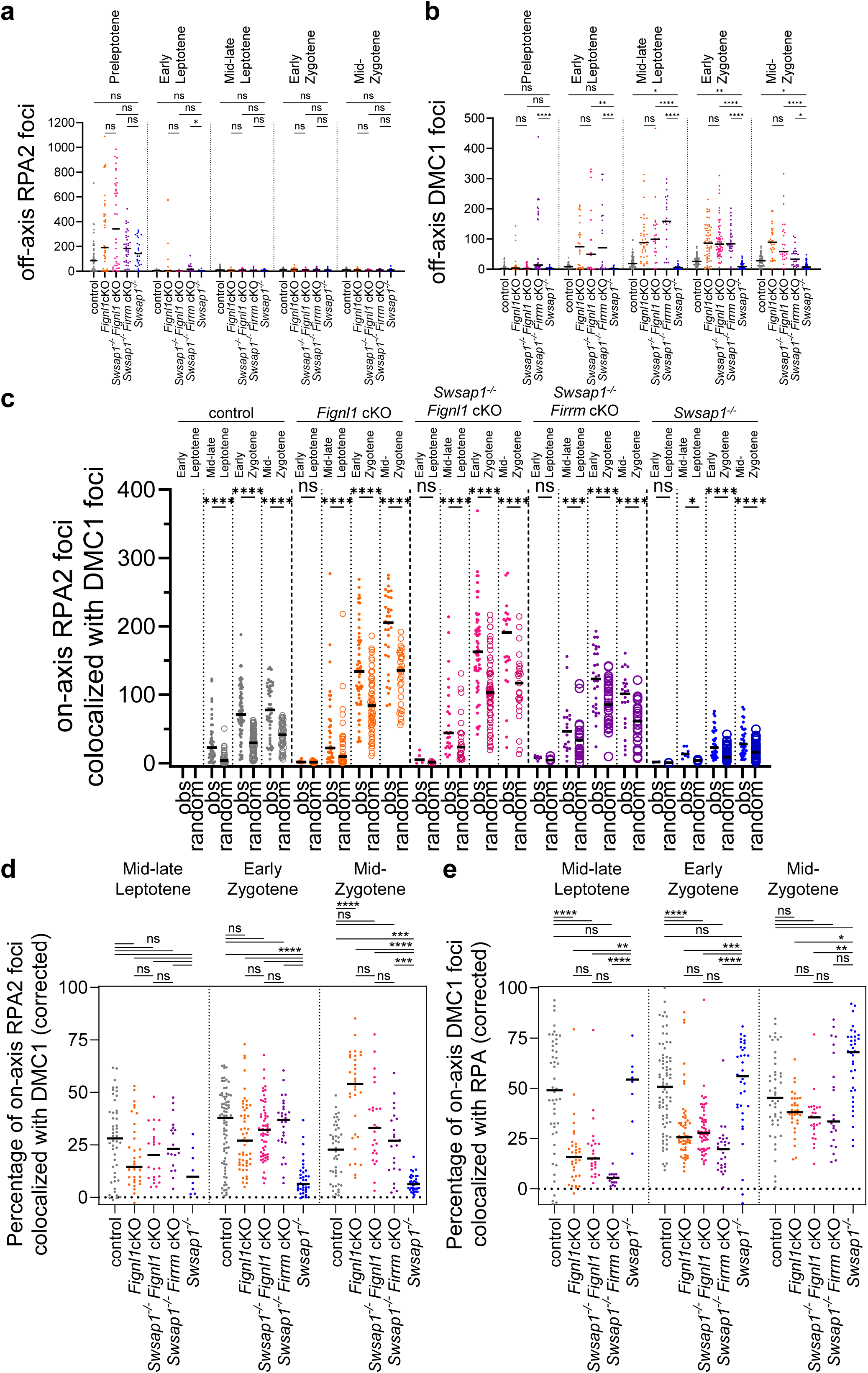
*Fignl1* deletion restores the formation of DMC1 foci in *Swsap1^-/-^* spermatocytes. **a-b.** Numbers of off-axis RPA2 **(a)** and DMC1 **(b)** foci detected on spermatocyte spreads from 17 dpp control mice of the indicated genotypes. n=1-3 mice per genotype (see legend of Fig. 7). **c-e.** Numbers **(c)** or percentages **(d,e)** of on-axis RPA2 foci colocalized with on-axis DMC1 foci **(c,d)** or vice-versa **(e)** in spermatocyte spreads from 17 dpp mice. The numbers of colocalized foci were corrected to the number expected by chance (see Methods).

**Extended Fig. 9.**
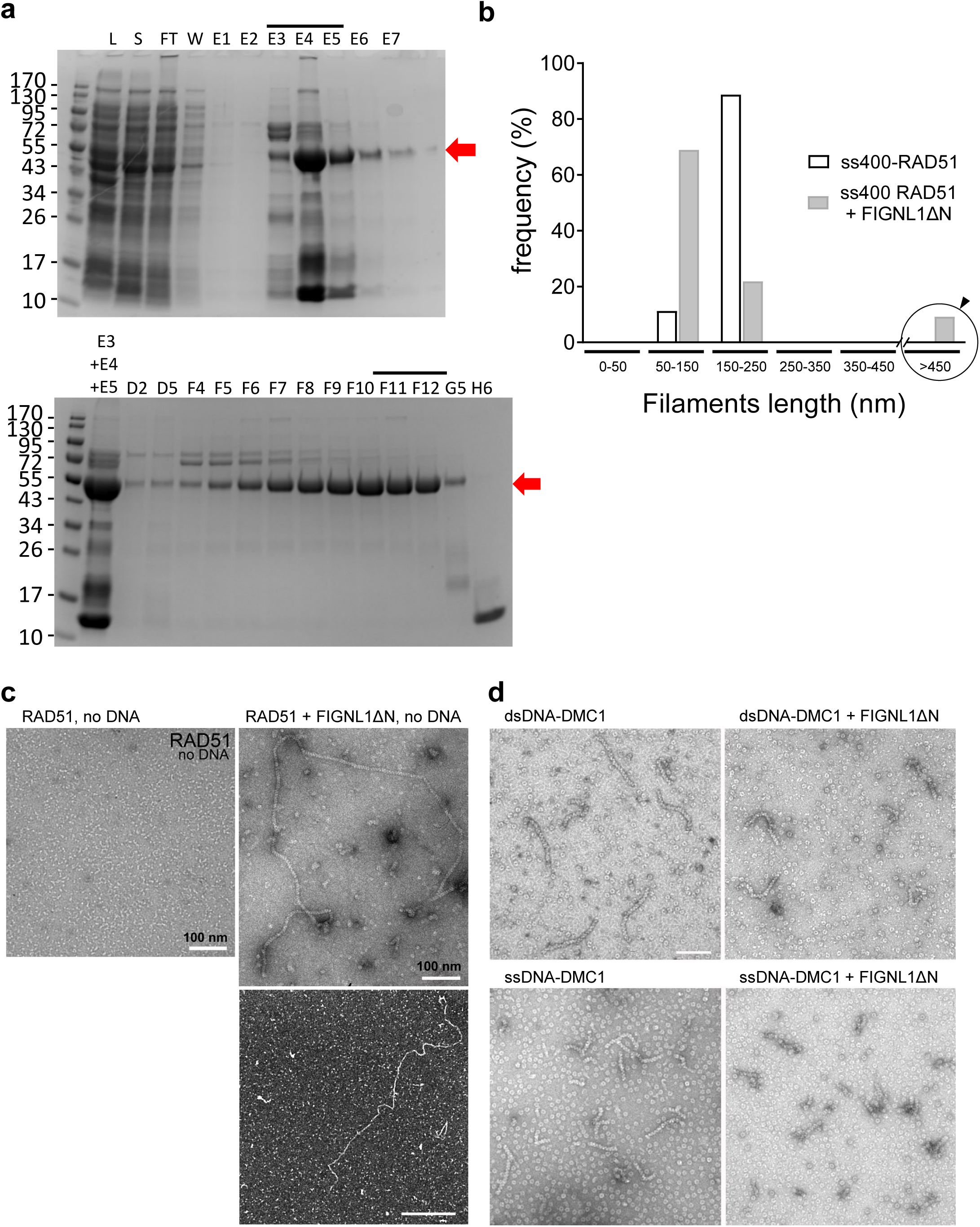
FIGLN1 alters the architecture and the activity of RAD51 and DMC1 nucleoprotein filaments. **a.** Purification of recombinant Histidine-tagged human FIGNL1ΔN284 protein from *E. coli*. Top panel, SDS-page analysis of proteins in total protein lysate (L), soluble protein fraction (S), flow-through (FT) from Hi-trap column, wash, and elution fractions (E1 to E7). Bottom panel, SDS-PAGE analysis of protein fractions collected during the gel filtration purification. Fractions E3, E4, and E5 from previous step were pooled and are shown as input control. Red arrows indicate recombinant His-FIGNL1ΔN284 with an expected size of 46kDa. F11 and F12 fractions were used for biochemical assays in this study. **b.** Length distribution of RAD51 filaments formed on 400 nt ssDNA fragments without (ss400-RAD51) or with (ss400-RAD51+ FIGNL1ΔN) 1.6 µM human FIGNL1ΔN. Note the presence of >450nm-long filaments when FIGNL1ΔN is present that were not included in the quantification shown in Figure 8f. **c.** Representative TEM images of RAD51 in the presence of ATP but in the absence of DNA (negative staining, left), and in presence of human FIGNL1ΔN (negative staining, scale bar 100nm, top right panel; and positive staining, scale bar 500nm, bottom panel). Note the presence of long filaments despite the absence of DNA. **d.** Representative TEM images (negative staining) of DMC1 filaments assembled on a 400 bp dsDNA (top) or 400 nt ssDNA (bottom) fragment, without (left) or with human FIGNL1ΔN (right). Scale bar, 100 nm.

**Source Data.**
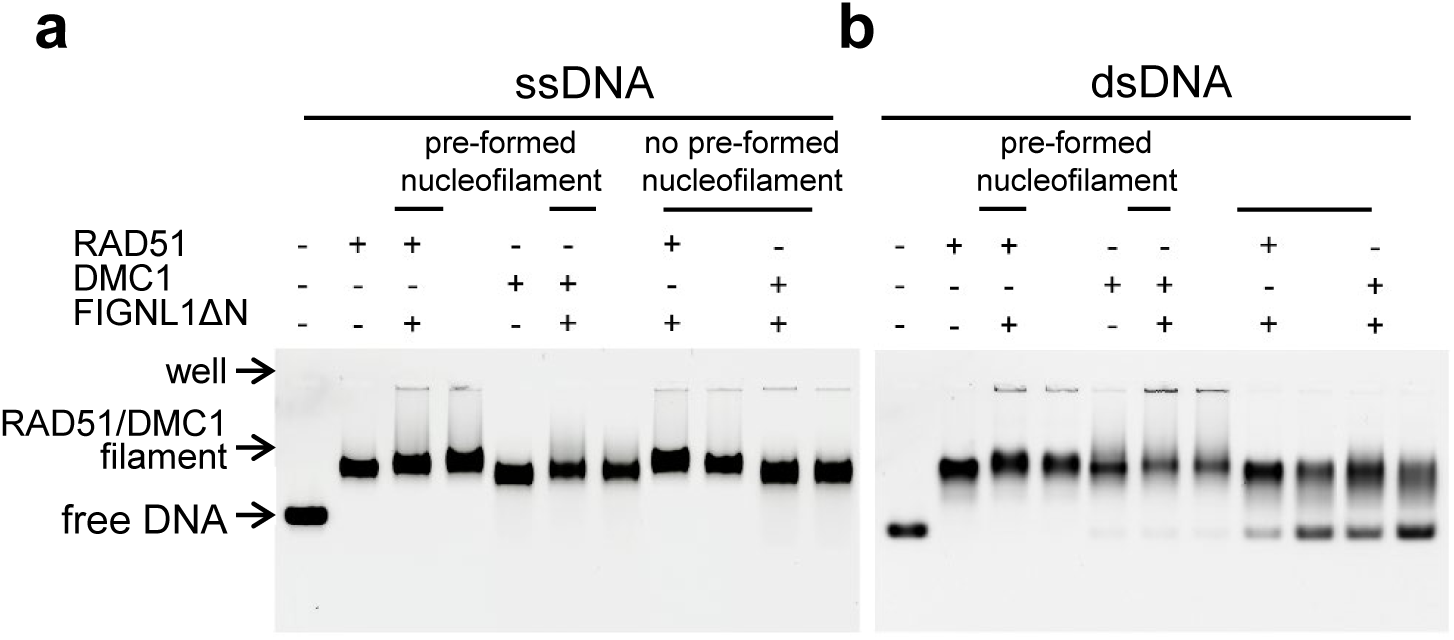
**a-b.** Uncropped image of the gels shown in Fig. 8a-b. Unlabeled lanes are not displayed on the final figure.

## Notes

### Competing Interest Statement

The authors have declared no competing interest.

### Summary of Updates

Additional analyses of Swsap1 KO Firrm cKO/Fignl1 cKO double mutants were included. We document the higher level of nuclear RAD51 measured by western blot in mutant testis nuclear extracts on one hand (Extended Data Fig. 2e), and the higher intensity of individual RAD51 (but not DMC1) foci (Extended Data Fig. 4a-b) on the other hand. Raphael Mercier was included as a co-author.

